# Purely STDP-based assembly dynamics: stability, learning, overlaps, drift and aging

**DOI:** 10.1101/2022.06.20.496825

**Authors:** Paul Manz, Raoul-Martin Memmesheimer

## Abstract

Memories may be encoded in the brain via strongly interconnected groups of neurons, called assemblies. The concept of Hebbian plasticity suggests that these assemblies are generated through synaptic plasticity, strengthening the recurrent connections within select groups of neurons that receive correlated stimulation. To remain stable in absence of such stimulation, the assemblies need to be self-reinforcing under the plasticity rule. Previous models of such assembly maintenance require additional mechanisms of fast homeostatic plasticity often with biologically implausible timescales. Here we provide a model of neuronal assembly generation and maintenance purely based on spike-timing-dependent plasticity (STDP) between excitatory neurons. It uses irregularly and stochastically spiking neurons and STDP that depresses connections of uncorrelated neurons. We find that assemblies do not grow beyond a certain size, because temporally imprecisely correlated spikes dominate the plasticity in large assemblies. Assemblies in the model can be learned or spontaneously emerge. The model allows for prominent, stable overlap structures between static assemblies. Further, assemblies can drift, particularly according to a novel, transient overlap-based mechanism. Finally the model indicates that assemblies grow in the aging brain, where connectivity decreases.

**Author summary:** It is widely assumed that memories are represented by ensembles of nerve cells that have strong interconnections with each other. It is to date not clear how such strongly interconnected nerve cell ensembles form, persist, change and age. Here we show that already a basic rule for activity-dependent synaptic strength plasticity can explain the learning or spontaneous formation and the stability of assemblies. In particular, it is not necessary to explicitly keep the overall total synaptic strength of a neuron nearly constant, a constraint that was incorporated in previous models in a manner inconsistent with current experimental knowledge. Furthermore, our model achieves the challenging task of stably maintaining many overlaps between assemblies and generating the experimentally observed drift of memory representations. Finally, the model predicts that if the number of synaptic connections in the brain decreases, as observed during aging, the size of the neuron ensembles underlying memories increases. This may render certain memories in the aging brain more robust and prominent but also less specific.

## Introduction

A widely used model of long-term memory posits that items are stored in the brain by strongly interconnected neuronal assemblies [1–3]. A memory item is represented by a group of neurons that coactivate upon memory recall. The assembly structure allows for associative recall from an incomplete input: due to the strong interconnections, activation of a part of the neurons in an assembly can trigger reactivation of the entire assembly and thus a recall of the full memory. The assemblies may each be created through experience-dependent plasticity or they may already form during development [4]. In the latter case, memories may form by connecting the pre-existing assemblies to appropriate input and output neurons [5]. The theory of the formation and maintenance of neuronal assemblies has been studied in much detail in previous works. The creation of new memories is commonly modeled using Hebbian plasticity [6–13]: if a set of neurons is co-activated, Hebbian plasticity increases the strength of their mutual connections leading to the formation of what is called a Hebbian assembly. However, memory networks in the brain also show ongoing spontaneous and irregular activity. If plasticity still takes place during this activity, it should not interfere with the existing memory assemblies – otherwise memories would have implausibly short lifespans. Hebbian assemblies can be self-reinforcing under plasticity since their interconnectedness leads to higher correlations in the activities, which in turn leads to potentiation of the intra-assembly weights. Models of assembly maintenance, however, found that fast homeostatic plasticity was needed in addition to Hebbian learning. This introduces competition between synapses and prevents pathological growth of assemblies and exploding activity [5, 7–9, 11–13]. Homeostatic plasticity has been observed in experiments, but it is much slower than Hebbian plasticity and does therefore not suffice to prevent runaway potentiation [14–18] (see, however, [19] for a different view and [10, 20] for a small timescale implementation of homeostasis via inhibitory STDP).

Experiments indicate that distinct memory assemblies have a fraction of shared neurons, i.e. neurons that are part of both assemblies [21–23]; the size of these overlaps appears to correspond to the strength of the associations between the concepts encoded by the assemblies. Previous models of assemblies stabilized by recurrent synaptic plasticity and fast homeostatic normalization usually do not show prominent overlaps [5, 7–9, 12]. An example of a network with weight plasticity, structural plasticity, multiple synapses per connection and short-term depression that can store two strongly overlapping assemblies was given in [24]. We will explore whether our purely STDP-based model can maintain overlaps.

Another topic of interest in the study of memory networks is whether they can generate the representational drift observed in recent experiments [25]. Such drift has recently been modeled by drifting assemblies, which spontaneously exchange neurons with each other, leading to a gradual resorting of the whole network [5]. The network model incorporated fast homeostatic normalization to stabilize the assemblies. We will explore whether also our purely STDP-based model can exhibit drifting assemblies.

Finally, aging and other conditions are associated with a decrease in overall connectivity [26, 27]. We will therefore explore how the assemblies in our networks adapt to such a decrease.

The paper is structured as follows: We initially introduce the model of spiking neurons and STDP and describe existing analytical approximations for the time-averaged weight dynamics. As a first result, we then show spontaneous assembly formation. We obtain an analytical approximation of the weight growth in different assembly sizes to obtain an understanding of the numerically observed assembly formation. Next we study whether assemblies can be learned by correlated external input. The subsequent section shows that our networks can stably maintain overlapping assemblies. We thereafter examine whether our model can be set up to exhibit representational drift. Finally, we investigate the dependence of assembly sizes on network sparsity and relate our results to effects of aging on the brain.

## Materials and methods

### Poisson neurons

Neural networks in the mammalian brain typically generate irregular and apparently largely stochastic spiking. To guarantee that our model networks generate similar spiking activity, we consider networks of stochastic linear Poisson (or “Hawkes”) model neurons [9, 29–31]. Such networks are additionally analytically well tractable. We explicitly model excitatory neurons. Since the irregular activity in biological neural networks is likely due to a balanced state, where excitatory and inhibitory inputs to each neuron balance [28, 32–36], our model implicitly incorporates inhibitory neurons, see Fig. 1.

**Fig 1.**
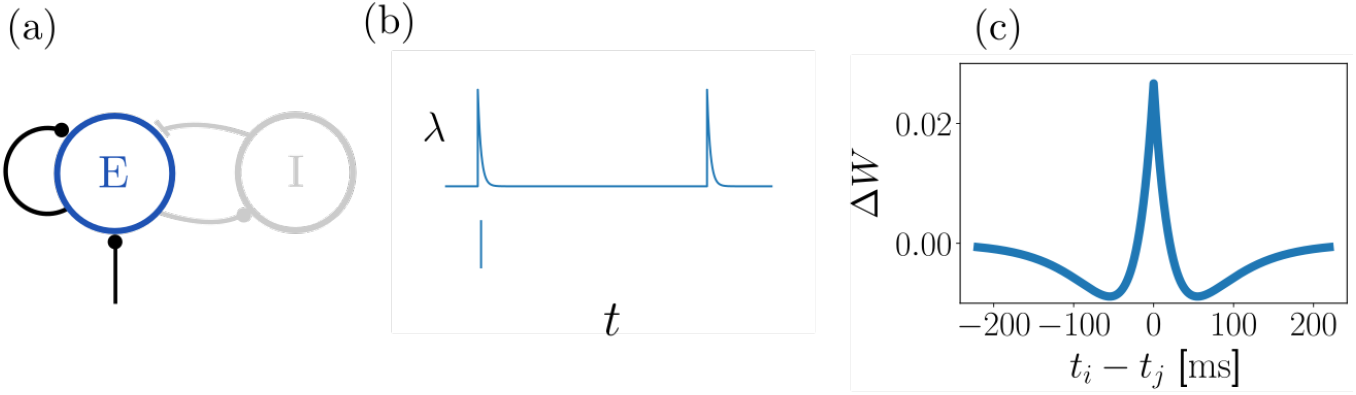
Network, neuron and plasticity model. (a): A network consists of recurrently connected excitatory neurons, which generate irregular, stochastic spiking. It may be interpreted as the excitatory population of a neural network where excitatory and inhibitory neurons generate a balanced state [28] of irregular spiking. Thus, a balancing inhibitory neuron population is implicitly contained in our model. (b): Poisson model of stochastic single neuron spiking: each neuron is characterized by its instantaneous rate *λ*(*t*) (upper subpanel), which depends on incoming spikes and determines the probability of emitting a spike (lower subpanel). (c): Strength of STDP updates as a function of the time difference between pre- and postsynaptic spikes (*μ* = 1).

The spiking activity of each neuron *i* is an inhomogeneous Poisson process whose time-dependent instantaneous spike rate (intensity) *λ*_*i*_(*t*) = ⟨*S*_*i*_(*t*)⟩ given input spike trains 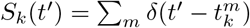 up to time *t* and weights *W*_*ik*_ is

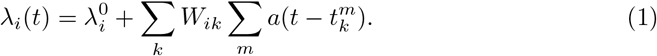

The angular brackets here denote trial-averaging with fixed input spike trains and weights to neuron *i*. We use an exponentially decaying synaptic kernel

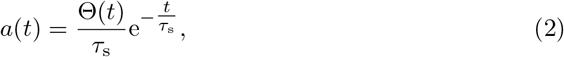

where Θ(*t*) is the Heaviside distribution, and a constant external drive *λ*^0^.

It is now useful to introduce quantities that are trial-averaged over the entire spiking network dynamics. The trial-averaged instantaneous rate (intensity) of neuron *i* is *r*_*i*_(*t*) = ⟨*λ*_*i*_(*t*)⟩, where angular brackets now denote the trial-averaging over the network dynamics. For static *W* with spectral radius less than 1, the trial-averaged rate dynamics relax to a fixed point where the vector *r*(*t*) is constant,

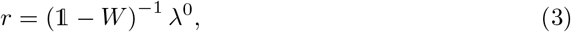

see [9, 30, 37, 38]. In addition, one can analytically compute the corresponding stationary cross-correlation functions *C*_*ij*_(*τ*) = ⟨*S*_*i*_(*t* + *τ*)*S*_*j*_(*t*)⟩ of pairs of neurons *i, j* at arbitrary *t*; in the frequency domain they read

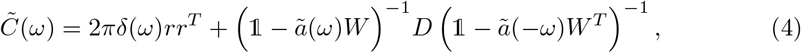

where we used matrix notation and *D*_*ij*_ = *δ*_*ij*_*r*_*i*_ [9, 30, 38]. *The Fourier transform of a function g*(*t*) is defined as 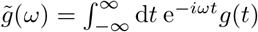. The first right hand side term in Eq. 4 is the correlation of independent Poisson spike trains with rate *r*, the second one reflects the interdependence of evoked spikes in the network. If the scale of the STDP updates is sufficiently small, one can assume that the weight dynamics are quasistationary with respect to neuronal dynamics. Then Eqs. 3,4 still approximately hold true despite *W* changing over time due to plasticity.

### Spike-timing-dependent plasticity

We consider networks of spiking neurons with pair-based STDP, i.e. the change of synaptic strength depends on the time lags of pairs of pre- and postsynaptic spikes. The characteristics of the plasticity function crucially determine how synaptic strengths evolve in a network. Networks with symmetric plasticity functions can establish a structure of neuronal assemblies whereas plasticity functions with a large antisymmetric part tend to establish feedforward chains of connectivity; see, however, [10, 20] for networks with asymmetric STDP maintaining assemblies and [12] for a triplet STDP rule that forms assemblies despite an asymmetric two-spike interaction part. In the present work, we consider symmetric plasticity functions because of their simplicity and analytical tractability, building on previous theoretical work that has employed them [5, 9]. Symmetric STDP has been found in the CA3 region of the rodent hippocampus [39], i.e. in a region that is assumed to serve as an associative memory network and to store assemblies [1]. Recently, near-symmetric STDP has also been observed in the primary motor cortex of primates [40].

The change induced by a spike pair with time lag *t* is given by the STDP function *F*,

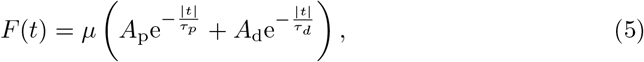

with *A*_p_ > 0, *A*_d_ < 0, |*A*_p_| > |*A*_d_| and *τ*_p_ < *τ*_d_, see Fig. 1c. We consider the overall scaling factor *μ* as the learning rate of the STDP function. *μ* is redundant and could be absorbed into *A*_p_ and *A*_d_, whose magnitudes also determine the scale of the STDP function. We keep *μ* because it facilitates the formulation of task conditions with the same STDP shape but different amplitudes. For the analytical treatment of our plasticity rule we set *μ* = 1. The assumption of small STDP scale to ensure a separation of timescales between plasticity and neuronal dynamics then becomes equivalent to assuming small *A*_p_ and *A*_d_. In all our simulations the parameters are chosen such that the integral 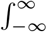 d*t F*(*t*) over *F* is negative.

At each spike time, plasticity acts additively on the pre- and postsynaptic weights of the spiking neuron, with amplitudes given by *F*. Specifically, at a spike time 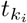 of the postsynaptic neuron there is a jump-like change in *W*_*ij*_ of 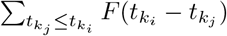. At a spike time 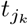 of the presynaptic neuron, *W*_*ij*_ jumps by 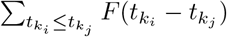. This can be compactly written as

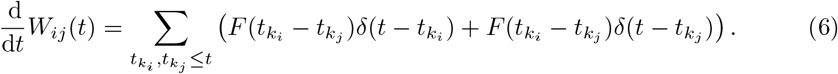

Here and henceforth we assume *i* ≠ *j*; there is no self-interaction in our networks, *W*_*ii*_ = 0. We further stipulate that no weight can become negative or exceed a maximum value *ŵ* due to STDP: if a synapse would be depressed below 0, it will be set to 0 instead; if a synapse would be potentiated to a value beyond *ŵ*, it will be set to *ŵ*.

In the regime of quasistationary weight dynamics the time-averaged drift in synaptic efficacy, 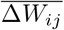, can be approximated by

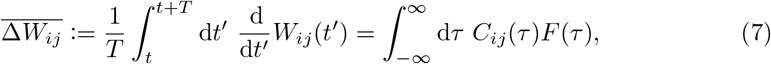

where *C*_*ij*_(*τ*) are again the correlations of the spike trains of neurons *i* and *j* [29]. We note that 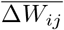 has the dimensionality of one over time. Plancherel’s theorem yields

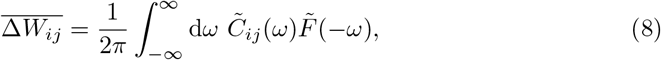

with the Fourier transforms of the correlation function and of the STDP window. Inserting the correlation function 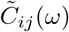 of Eq. 4 gives

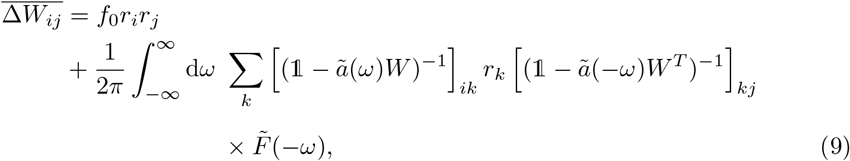

see [9], where

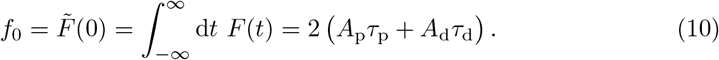

It is often useful to expand Eq. 4 into a power series with respect to *W* [9, 30]. Inserting the series into the right hand side of Eq. 8 (or directly expanding Eq. 9) results in a series expansion for 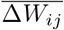, [9]:

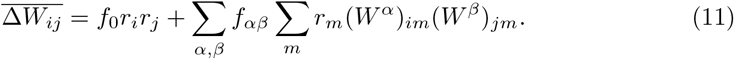

The terms of the sum encode contributions from motifs in which a source neuron affects post- and presynaptic neurons via a chain of *α* and *β* connections, respectively (the same connection may be counted more than once). If *α* = 0 (*β* = 0) the post(pre)synaptic neuron itself is the source. The coefficients *f*_*αβ*_ contain integrals over the Fourier transform of the STDP window and powers of the Fourier transform of the synaptic kernel function,

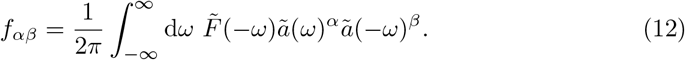

### Simulation parameters

We have used the following parameters in the network simulations for the figures below.

Fig. 2: The network starts with an initially random connectivity where each weight is independently and uniformly drawn from [0, 0.25*ŵ*].

**Fig 2.**
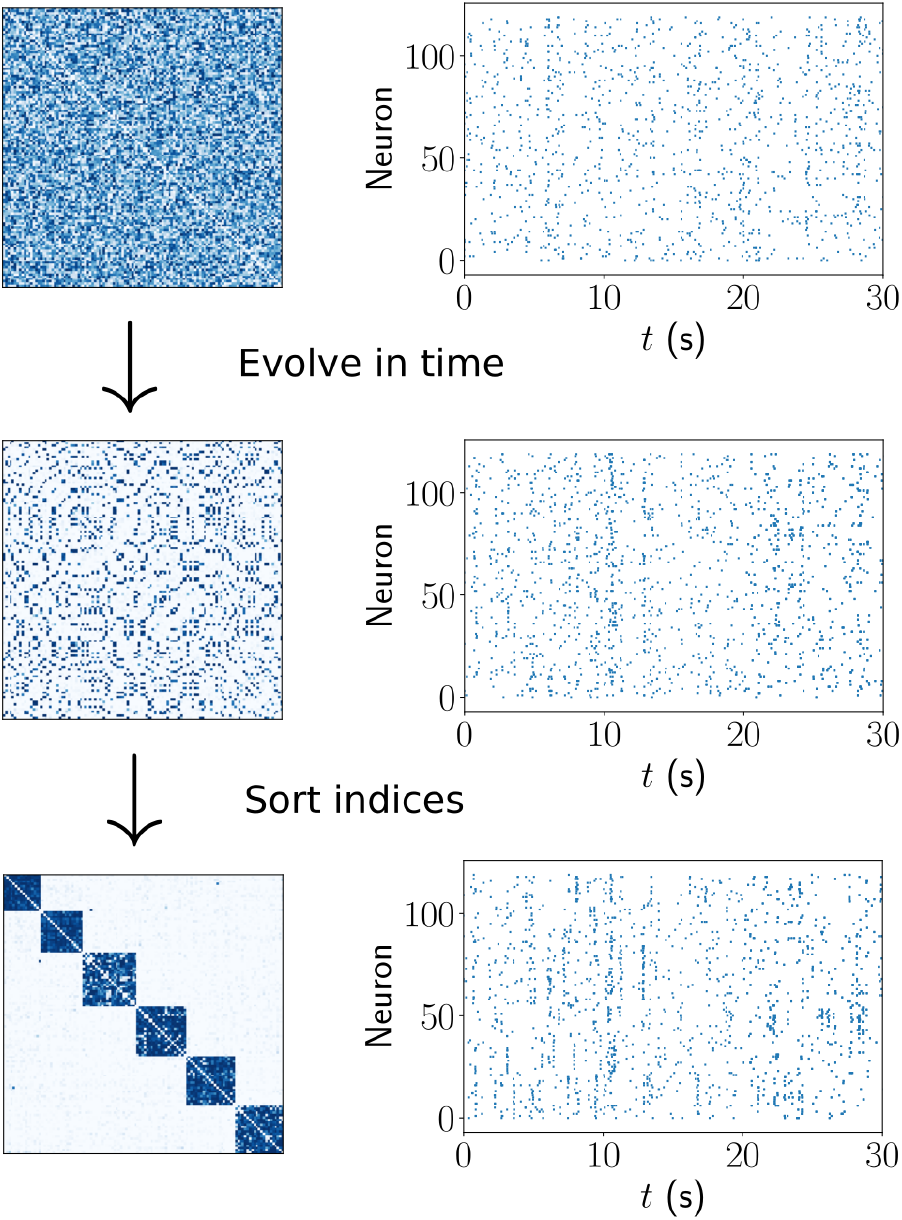
Spontaneous assembly formation. Several assemblies (strongly interconnected ensembles of neurons) emerge in a network with initially random connectivity, due to spontaneous activity.

Fig. 3b: The networks start with an initially random connectivity where each weight is independently and uniformly drawn from [0, 0.20*ŵ*]. We simulate for 10 different initial conditions and then take the median of all final assemblies.

**Fig 3.**
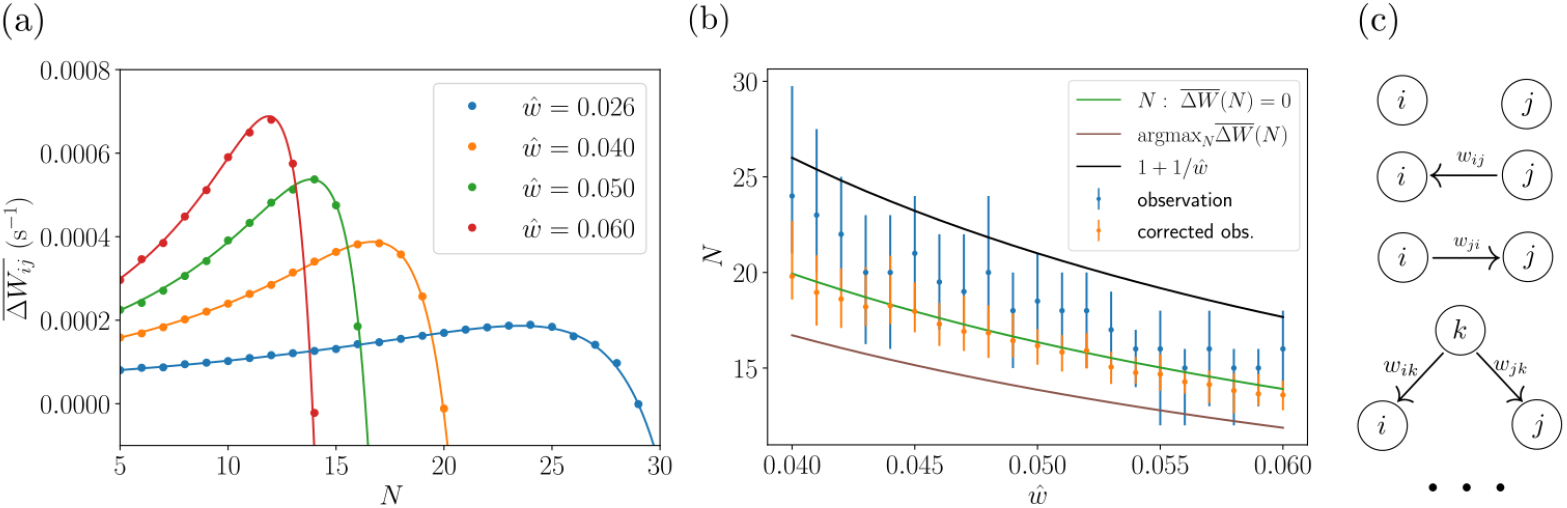
Explanation of assembly formation. (a): Time-averaged weight change (tracked, not applied) for a synapse in an assembly (fully connected, no other connections, all weights fixed at *ŵ*) as a function of its size *N*, for different values of *ŵ*. Solid lines: theoretical prediction, dots: numerical results from fixed homogeneous assemblies. (b): Sizes of spontaneously forming assemblies in network simulations with different *ŵ* (blue dots: medians, error bars: first and third quartiles) and corresponding sizes of assemblies with homogeneous coupling of strength *ŵ* computed using a correction for sparseness (orange). The lower continuous curves show analytical predictions for the sizes of homogeneously connected assemblies obtained from Eq. 15, namely the solutions of 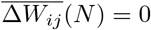 (green) and argmax 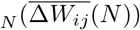 (brown). The upper curve displays the analytical upper limit of the homogeneously connected assembly sizes, 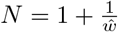 (black). (c): Illustration of zeroth, first and second order contributions to the time-averaged weight change for a given rate and weight configuration. Circles and arrows represent neurons and synaptic connections, respectively.

Fig. 4: The initial weight matrix has an assembly of neurons 1-20 interconnected with *W*_*ij*_ = *ŵ* and background connectivity as in Fig. 2. During the stimulation period neurons 21-40 are given input from a single source of Poisson spiking with *λ* = 4.18 Hz and input weights *w*_in_ = 10*ŵ*.

**Fig 4.**
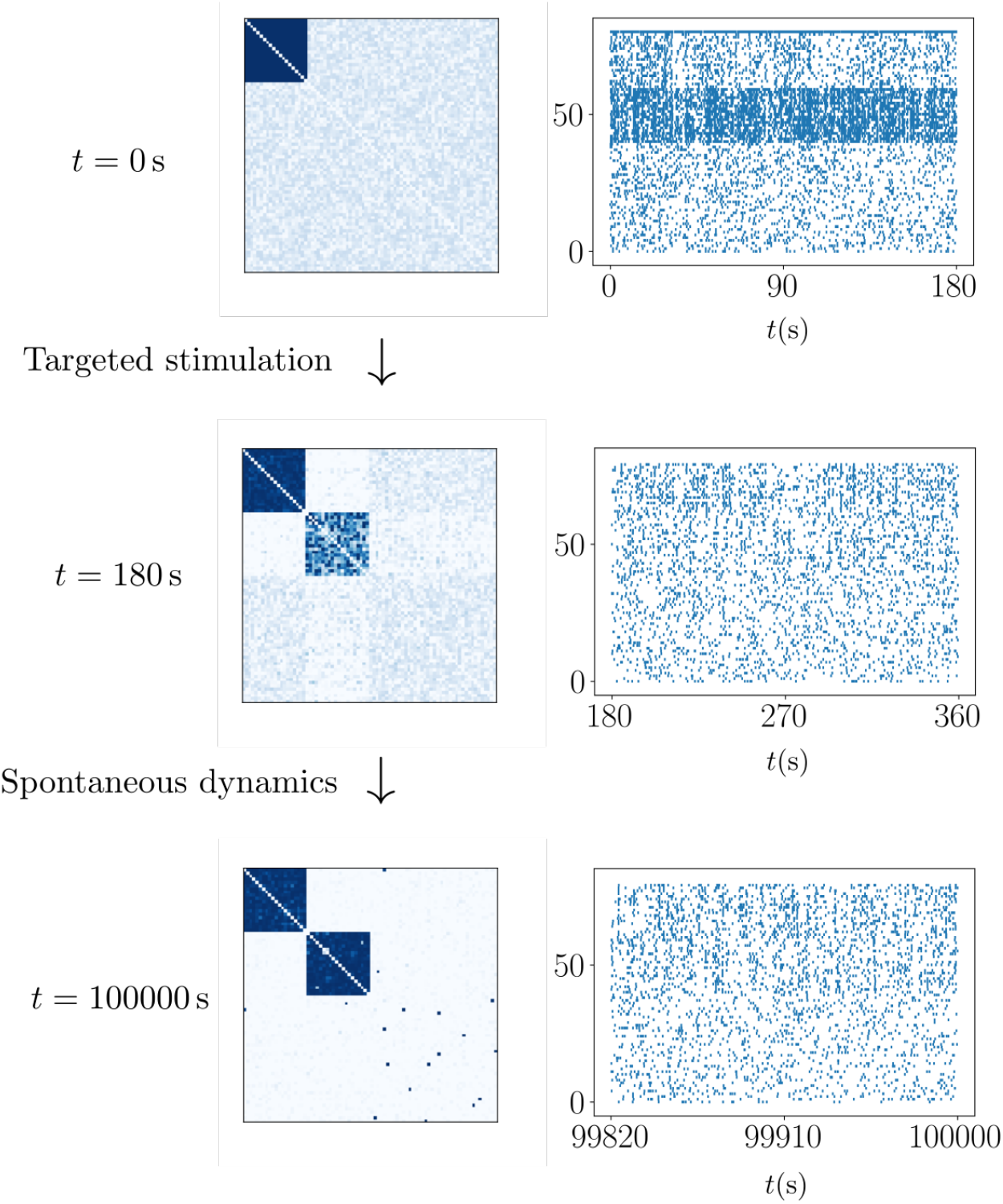
Learning a new assembly in a network starting with a single assembly and otherwise weak random connectivity. Initially, correlated external input stimulates the neurons of the new assembly, which leads to an increase of their interconnecting weights and thereby to the formation of a rudimentary assembly. The stimulation is then turned off and the network evolves on its own; the new assembly becomes fully connected.

Fig. 6a: The neuron with index 10 has spontaneous rate *λ*_0_ = 0.08 Hz.

Fig. 6b: The neuron with index 10 has spontaneous rate *λ*_0_ = 0.01 Hz.

Fig. 8a: The initial weight matrix is as in Fig. 2 but with weights drawn from [0, 0.15*ŵ*].

Fig. 8b: Every 3 × 10^5^ s a neuron changes its spontaneous rates with probability *p* = 0.03 to *λ*^*′*^_0_ = 0.03 Hz for 3 × 10^5^ s. For the last 3 × 10^5^ s of the simulation all spontaneous rates are kept at *λ*_0_ = 0.15 Hz.

## Results

### Spontaneous assembly formation

We first simulate the model described above with initially unstructured weight matrix and without structured external stimulation, in order to test its capability of spontaneous assembly formation and subsequent maintenance. We find that for appropriately chosen parameters in the plasticity function the network weights indeed converge towards a structure with segregated assemblies of a characteristic size.

The mechanism underlying the increase of weights between future assembly neurons is well known: Initially basically randomly coincident spiking leads to strengthening of some weights. By directly and indirectly coupling neurons, these weights induce more near-simultaneous spiking, which further strengthens them. The positive feedback loop leads to weights that increase until they reach *ŵ* [41]. Neurons with little coupling do not display similar near-simultaneous spiking such that their weights grow more slowly, or even shrink due to long-term depression. It has been shown that if the summed weight strength to and from a neuron are additionally normalized by fast homeostasis, there can be spontaneous emergence of assemblies [5, 9, 12, 13]. The reason is that the homeostatic normalization lets synapses compete, such that more slowly growing ones are suppressed. Once an imbalance of connectivity and thus first assembly-like structures occur, the weight increase of intra-assembly synapses prior to normalization is stronger, due to the stronger co-spiking of the connected neurons. The normalization then suppresses the inter-assembly connections and the assembly structure consolidates.

While the activity-dependent plasticity-based mechanisms underlying spontaneous assembly formation remain in our networks, there is no fast homeostatic normalization that could introduce competition between synapses. Then, why do not all weights tend to *ŵ*? In other words: why does the network not turn into one big assembly?

### Plasticity in homogeneously connected assemblies

In this section, we will argue that the assemblies in our model grow if their size is small and that their final size is limited. We explain the latter by the depression dominance of the learning rule and imprecisely correlated spiking dominating the plasticity in large assemblies. To model assembly weight dynamics, we consider the special case of an isolated assembly of *N* neurons that is homogeneously connected with synaptic strengths *ŵ*. We compute the time-averaged weight change

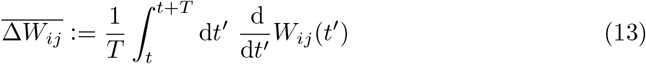

(cf. Eq. 7) for weights within this type of assembly, disregarding the clipping at *ŵ*. This indicates whether and how vigorously weights that fall below *ŵ* are restored towards it. Further, it indicates how successfully further neurons are recruited, as recruitment relies on increasing the weights to new neurons once they are randomly increased. We expect that the assembly size is limited by values of *N* above which 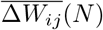 is negative: Larger assemblies will lose neurons due to the depression of weights, to other, smaller assemblies that tend to potentiate their connections with these neurons. This yields as an estimate for the size *N* of assemblies the solution of 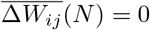. The assumption that an assembly loses neurons already if another assembly can potentiate its connections to them more yields as preferred assembly size the *N* that maximizes 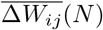.

To analytically obtain 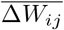 as a function of the assembly size, we first compute from Eq. 3 the average rate of an assembly neuron

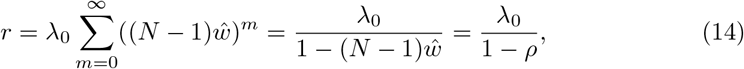

where *ρ* = (*N* − 1)*ŵ* is the branching parameter, which gives the average number of spikes that are directly induced by a single spike in the assembly. *N* is restricted to 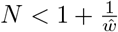 because for larger values there is no stationary network state with finite firing rates. This is reflected by the divergence of the geometric series in Eq. 14. In other words, a stable network requires *ρ* < 1 [31, 42, 43]. Eqs. 4, 14 and 8 yield

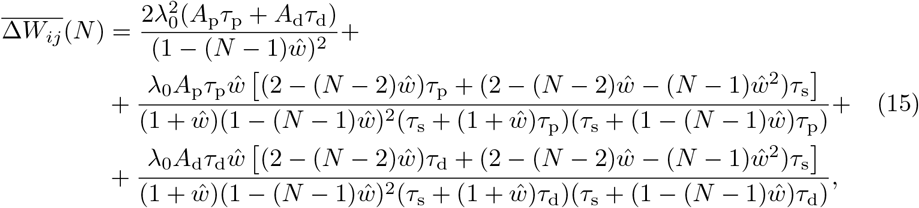

see SI S.1 for details. The first right hand side term covers the impact of uncorrelated pre- and postsynaptic spiking with rates *r*_*j*_ and *r*_*i*_ on the synaptic strengths (*C*_*ij*_(*τ*) = *r*_*i*_*r*_*j*_ for uncorrelated spiking). The second and third terms, which are proportional to *A*_p_ and *A*_d_, respectively, cover the weight potentiation and depression due to correlated spiking.

We now study under which conditions the weights of small assemblies tend to increase. For this we assume small *N* ≥ 2, such that together with *ŵ* ≪ 1, which holds for the parameters that we employ throughout our article, also *N ŵ* ≪ 1. Eq. 15 then simplifies to

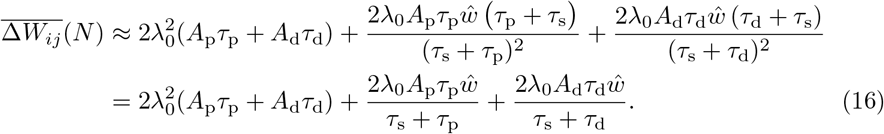

We require the right hand side to be larger than zero. To interpret this criterion, we rewrite it in terms of the integrals of the potentiating part 2*A*_p_*τ*_p_ and of the depressing part 2*A*_d_*τ*_d_ of the STDP window and multiply by *τ*_s_,

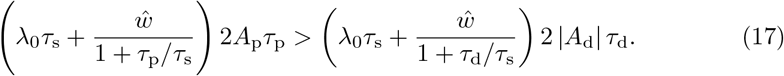

The integrals 2*A*_p_*τ*_p_ and 2*A*_d_*τ*_d_ characterize the overall strengths of potentiation and depression. *λ*_0_*τ*_s_ and *ŵ* weight the strength of spontaneous, always uncorrelated spiking within a synaptic time constant against the induced spiking by an input spike.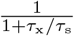 describes how effectively the time constants allow induced spikes to translate to plasticity: Since we have *ŵ* ≪ 1 and small *N*, low order connectivity motifs are most relevant for spike generation. These generate pre- and postsynaptic spikes with a small temporal delay on the order of *τ*_s_. The evoked potentiation or depression is most effective if 2*A*_x_*τ*_x_ is concentrated to a narrow peak, i.e. *τ*_x_ is small. The effect saturates if *τ*_x_ falls below *τ*_s_, because *τ*_s_ limits the precision of evoked correlated spiking.

For the parameters used in our paper, Eq. 17 is satisfied. It does not hold, however, for overly small *ŵ*, see our discussion in SI S.1 and Fig. S3. For the parameters used in our paper, although the overall strength of depression is larger than that of potentiation, the narrower peak of the latter (*τ*_*p*_ < *τ*_*d*_) implies that potentiation will be more efficiently triggered by short-range spike correlations and therefore dominate in small networks where firing rates are lower and lower order connectivity motifs are more important.

To show that the size of assemblies is limited, we neglect *N*’s discrete nature and consider the asymptotic behavior of 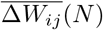 with *N* approaching 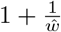. For this we insert 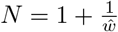 in Eq. 15 except in the factor that leads to divergence,

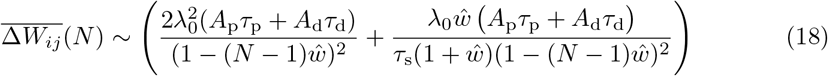

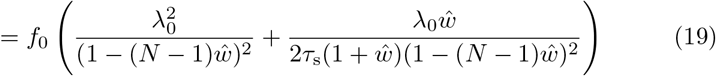

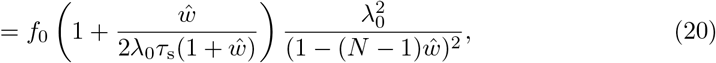

where we used Eq. 10 to obtain the second line. The result implies

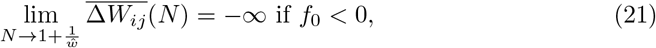

since the summands in the large bracket and the trailing factor are positive. Thus, if *f*_0_ < 0 and there is an *N* for which 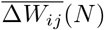 is positive, 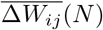 will have a positive maximum and a zero at an *N* smaller than 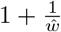 (cf. also the phase diagram Fig. S3); we therefore expect limited assembly growth and assume *f*_0_ < 0 throughout the article. In contrast, if 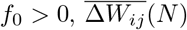 diverges to positive values for *N* where also the firing rate diverges, indicating that sufficiently large assemblies continue to grow until the network generates pathological activity.

The first terms in the large brackets of Eqs. 18-20 cover the impact of uncorrelated pre- and postsynaptic spiking. As the firing rates increase with *N* and *f*_0_ < 0, the contribution of the STDP rule due to uncorrelated spiking tends to increasingly negative values. The fact that the impact of the second term, which encodes the effects of connectivity motifs on STDP (Eq. 11), also becomes negative for sufficiently large *N* is due to contributions from higher order motifs. We can see this by reconsidering

Eq. 11: For homogeneously connected assemblies, it simplifies to

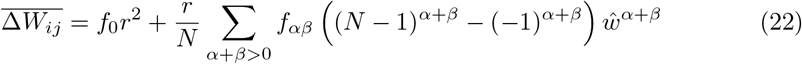

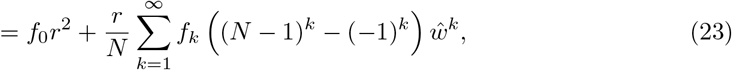

where

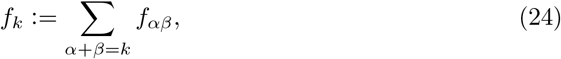

see SI S.2. Since *f*_*αβ*_ (cf. Eq. 12) and thus *f*_*k*_ are independent of *N*, higher order terms in *k* grow in Eq. 23 faster with *N* than lower order ones. Therefore higher order motifs become more relevant for the weight change the larger *N* is. We find for high orders that

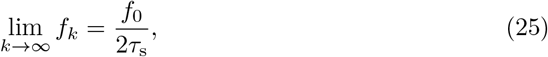

see SI S.2 and Fig. S1; therefore high order motifs induce synaptic depression. This depression is impactful: For our sets of parameters, *λ*_0_*τ*_s_, the average number of spontaneous spikes generated within the duration of a synaptic current, is considerably smaller than 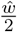, half the average number of spikes generated by an incoming spike at a neuron (see Table 1). Further, *ŵ* is considerably smaller than 1. The large bracket in Eq. 20 thus tells us that the imprecisely correlated spiking due to higher order motifs and not the overall increase in spike rate in the assemblies is mostly responsible for the decrease of 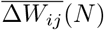 for 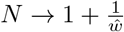.

**Table 1.** List of parameters for the simulations in the referenced figures

| | $N$ | $\hat{w}$ | $\tau_s$ | $A_p$ | $A_d$ | $\tau_p$ | $\tau_d$ | $\mu$ | $\lambda_0$ |
| --- | --- | --- | --- | --- | --- | --- | --- | --- | --- |
| Fig. 1c |  |  |  | 0.08 | -0.0533 | 0.025 s | 0.05 s |  |  |
| Fig. 2 | 120 | 0.04 | 0.01 s | 0.08 | -0.0533 | 0.025 s | 0.05 s | 0.07 | 0.15 Hz |
| Fig. 3a |  |  | 0.01 s | 0.08 | -0.0533 | 0.025 s | 0.05 s |  | 0.15 Hz |
| Fig. 3b | 150 |  | 0.01 s | 0.08 | -0.0533 | 0.025 s | 0.05 s | 0.04 | 0.15 Hz |
| Fig. 4 | 80 | 0.026 | 0.01 s | 0.08 | -0.066 | 0.035 s | 0.05 s | 0.07 | 0.15 Hz |
| Fig. 5b | 120 | 0.017 | 0.01 s | 0.08 | -0.044 | 0.022 s | 0.055 s | 0.02 | 0.15 Hz |
| Fig. 6a | 60 | 0.0158 | 0.01 s | 0.08 | -0.044 | 0.022 s | 0.055 s | 0.021 | 0.15 Hz |
| Fig. 6b | 60 | 0.024 | 0.01 s | 0.08 | -0.0533 | 0.025 s | 0.05 s | 0.045 | 0.15 Hz |
| Fig. 7 | 190 | 0.018 | 0.01 s | 0.08 | -0.042 | 0.026 s | 0.065 s | 0.017 | 0.15 Hz |
| Fig. 8a | 72 | 0.056 | 0.01 s | 0.08 | -0.0533 | 0.025 s | 0.05 s |  | 0.2 Hz |
| Fig. 8b | 120 | 0.024 | 0.01 s | 0.08 | -0.0533 | 0.025 s | 0.05 s | 0.05 | 0.15 Hz |
| Fig. 9a |  | 0.058 | 0.01 s | 0.08 | -0.0533 | 0.025 s | 0.05 s |  | 0.15 Hz |
| Fig. 9b | 120 | 0.058 | 0.01 s | 0.08 | -0.0533 | 0.025 s | 0.05 s | 0.07 | 0.15 Hz |
| Fig. 9c | 150 | 0.067 | 0.01 s | 0.08 | -0.0533 | 0.025 s | 0.05 s |  | 0.2 Hz |

Importantly, the relation Eq. 25 holds generally, for any plasticity window *F*, not only for the considered symmetric one. We explain it as follows: A term with specific exponents *α* and *β* in Eqs. 11, 22 covers the contribution of a specific connectivity motif to the spike correlation. This motif consists of a common presynaptic neuron that is separated from neurons *j* and *i* by a chain of *β* and *α* connections, respectively [9], see Fig. 3c for an illustration. Higher orders of *k* = *α* + *β* thus encode the effects of long cascades of spiking activity in an assembly. The longer these cascades get, the more spread-out and Gaussian the temporal distribution of their impacts becomes, due to the summation of inter-spike-intervals and their jitters, see Fig. S1. (We note that since the considered model is linear, one can indeed attribute the generation of each spike to a precursor spike or to the external drive.) Eq. 24 homogeneously sums over motifs with different connection chain lengths to the pre- and the postsynaptic neuron. For sufficiently large *k* the evoked spike time differences are thus equidistant, broad, overlapping Gaussians over the STDP window, which leads to the dependence on the integral of the STDP function *f*_0_, as for uncorrelated pre- and postsynaptic spiking.

The broadening of temporal spike correlations for 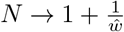 and its impact on 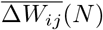 can also be observed from the interdependence term of 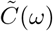 (cf. Eq. 4) without expanding it. SI S.2 (last paragraph) shows that it tends in the time domain to a two-sided exponential with smaller and smaller decay time constant. In the limit there is no more decay; the correlation function becomes flat. This implies together with the flat time domain correlation function of Poisson spike trains (cf. the first term of Eq. 4) and Eq. 7 that the weight change 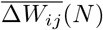 only depends on the integral *f*_0_ of *F*.

Fig. 3a shows 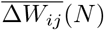 as in Eq. 15 for different values of 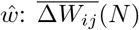 is maximal for a particular assembly size, then crosses zero and becomes increasingly negative. We confirmed these results using simulations of homogeneous assemblies, in which the weight updates that would occur due to STDP without clipping are tracked but not applied and then averaged over time. In Fig. 3b we show the typical assembly sizes that emerge for different values of *ŵ* in network simulations of spontaneous assembly formation. We find dense but also sparser assemblies where some internal connections are weak or basically missing, see, for instance, the third assembly in Fig. 2. For these we expect the maxima and zero crossings of 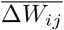 to be at larger sizes compared to our estimates with fully connected assemblies. Consistent with this, we observe that sparser assemblies in our simulations tend to be larger. In Fig. 3b, we correct for the effect of missing connections by taking the sum of the intra-assembly connections of an assembly, *w*_sum_, and computing the size *N*_corrected_ that a homogeneously connected assembly with the same *w*_sum_ and weights *ŵ* would have,

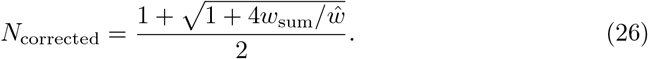

The corrected assembly sizes closely match the ones predicted from the zeros of 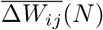, while the prediction from the maxima of 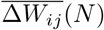 is farther off. This indicates that the assemblies grow until further growth would result in a decrease of the intra-assembly weights. The corrected assembly sizes lie below and scale differently than the upper limit for fully connected assemblies, 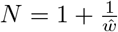. This is consistent with the theoretical prediction that assemblies reaching this size would quickly lose intra-assembly weight because 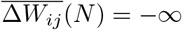.

The positive part of the 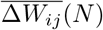 curve shows that for our parameters the potentiating STDP part dominates for small and intermediate *N*. It reflects the spontaneous formation of assemblies due to precise spike correlations as in previous work and their tendency to grow bigger for small *N*, see SI S.1. We note that Eq. 15 may suggest to take the limit of *ŵ* → 0 in order to simplify it, or to assume that the dynamical quantity *N* goes to infinity, which requires that the system parameter *ŵ* becomes infinitesimally small to keep the rates finite for all admissible *N* (cf. Eq. 14). *ŵ* → 0, however, implies that 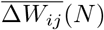 becomes negative for all *N*, since the rate-rate-interaction term dominates, see SI S.1, last paragraph, and Fig. S3. Therefore our theory predicts that no assemblies form in this limit.

If we truncate our theory to correlations evoked by direct weight connections and common presynaptic neurons [44] (i.e. to the terms *f*_0_, *f*_10_, *f*_01_, *f*_11_ in Eq. 11), 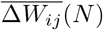 also has a maximum, crosses zero and then tends to negative infinity. This decrease is a purely rate-based effect: it occurs because the magnitude of the rate-rate interaction term proportional to *f*_0_ increases quadratically with the firing rate (the other terms are positive and linear in the rate). We compare two examples of 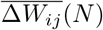 from the truncated theory with their full theory counterparts in Fig. S2. For the larger value of *ŵ*, the maximum and zero of the truncated theory curve occur at higher values of *N* compared to the full theory. In contrast, leaving out the rate-based term *f*_0_*r*_*i*_*r*_*j*_ in Eq. 11 has only little impact. For the smaller value of *ŵ* both modifications lead to shifts that are significant and similar; the curve without rate based term starts to drop faster than the truncated one only beyond the zero. These observations show the importance of imprecisely correlated spiking for limiting the assembly growth through the higher order motif- and the rate-based term in Eq. 11. They are consistent with the dominance of the motif-based term in the asymptotics Eq. 20, which is more pronounced for larger *ŵ*.

We conclude that the basic mechanism of spontaneous assembly formation due to activity-dependent plasticity, which we described in the previous section, remains in our model. For the limitation of assembly size, however, we uncover a novel mechanism: temporally imprecisely correlated spikes dominate the plasticity in large assemblies and do not allow them to grow further. The imprecision results from the long pathways between a spike and the spikes that it indirectly generates. The depressing effect of the imprecisely correlated spikes is covered by the negative rate-rate interaction term and by the negative total contribution of higher order motifs; both are proportional to *f*_0_. Eq. 15 includes their impact and quantitatively describes how the size of spontaneously emerging assemblies remains limited in our model.

### Storing new assemblies

Next we show how assemblies can be learned via correlated feedforward input. Neurons recruited for a new assembly may previously have only weak connections, i.e. they may belong to a background of neurons in front of which assemblies exist. Alternatively recruited neurons may already be part of other assemblies. In our simulations, during the learning phase, each neuron that is to be recruited receives Poisson spike input from the same source. This stimulates the neurons to spike near-simultaneously, such that the weights between them grow. Fig. 4 displays the resulting weight and spiking dynamics in a network which prior to stimulation hosts one assembly in front of a background of weakly coupled neurons. Before time *t* = 0s, the single assembly is stably stored in the network, despite the ongoing plasticity. At *t* = 0 s a group of background neurons receives correlated stimulation for *T*_stim_ = 180 s. This leads to the formation of a rudimentary assembly. After the stimulation has ended, its synapses grow over a longer period of time until a fully connected assembly is reached, where all interconnections have synaptic strength close to *ŵ*. The remaining background neurons, which did not receive correlated stimulation in the beginning, do not form assemblies. The stimulation furthermore does not affect the already existing assembly and neither does this assembly interfere with the formation of the new assembly. Fig. S6 shows another example of assembly learning; there again mostly background neurons are recruited, but also one neuron that is already part of two pre-existing assemblies.

Assembly creation requires that there is net potentiation 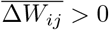 of the weights between the stimulated neurons. Using this condition and assuming that initially the weights between assembly neurons are zero, Eq. 11 yields an approximate analytical lower bound for the required strength of the external input given the input rate, see SI S.3 for details. Numerical simulations confirm this bound and show that noise allows assemblies to form already for smaller weights, see Fig. S4.

### Overlapping Assemblies

Experiments indicate that neurons can code for more than one memory item [21–23]. For assembly models of memory this implies that neurons can be part of more than one assembly, see Fig. 5a, i.e. assemblies have overlaps [45]. Such overlaps may encode associations between memories.

**Fig 5.**
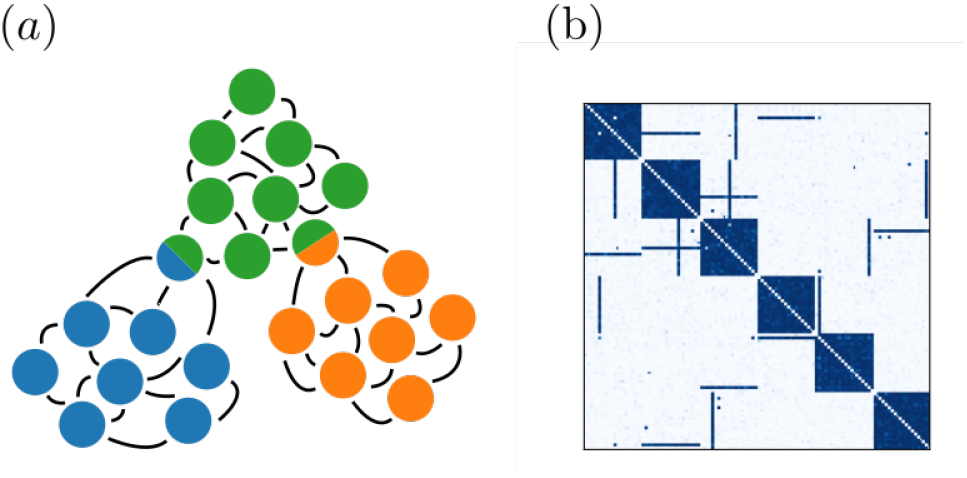
Overlapping assemblies. (a): Illustration of overlaps. (b): Weight matrix of a network with overlapping assemblies. The base assemblies (diagonal blocks) are each 20 neurons in size.

We find that for appropriate parameters, our networks can stably store assemblies with some overlaps, see Fig. 5b for an example weight matrix. However, we also observe that overlaps present a challenge for our models: overlapping assemblies either tend to merge or overlap neurons tend to disconnect from one of the assemblies they are a part of, dissolving the overlap. The first case occurred especially when the sizes of the overlapping assemblies were significantly smaller than the sizes that would lead to 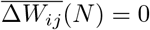. We hypothesize that in this case a feedback loop arises: overlaps induce correlations between the assemblies, which facilitates the formation of additional overlaps until the assemblies have completely fused. The other case occurred when the sizes of the two assemblies were close to 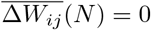. We hypothesize that here the higher firing rate of an overlap neuron and the resulting increased synaptic depression imply that overlaps become more likely to disappear due to fluctuations in the weight dynamics.

The hypotheses about the mechanism underlying the problems with overlap suggest two solutions how networks may solve them: First, if the synaptic long-term depression that occurs at high rates limits the connectivity of neurons, a neuron with a significantly lower spontaneous rate may be able to connect to multiple assemblies at the same time. In Fig. 6 we show how neurons with lower spontaneous firing rate join a second assembly. In panel (a) a neuron with lower spontaneous rate is initially already partially connected to another assembly. This partial connection then causes enough correlation with the second assembly to induce the completion of the connections. In this case both the initial partial connection and the lower *λ*_0_ were necessary to create this overlap. For an even lower *λ*_0_, an overlap emerged spontaneously, with a randomly selected assembly, through the inherent stochasticity of the dynamics, as shown in Fig. 6b.

**Fig 6.**
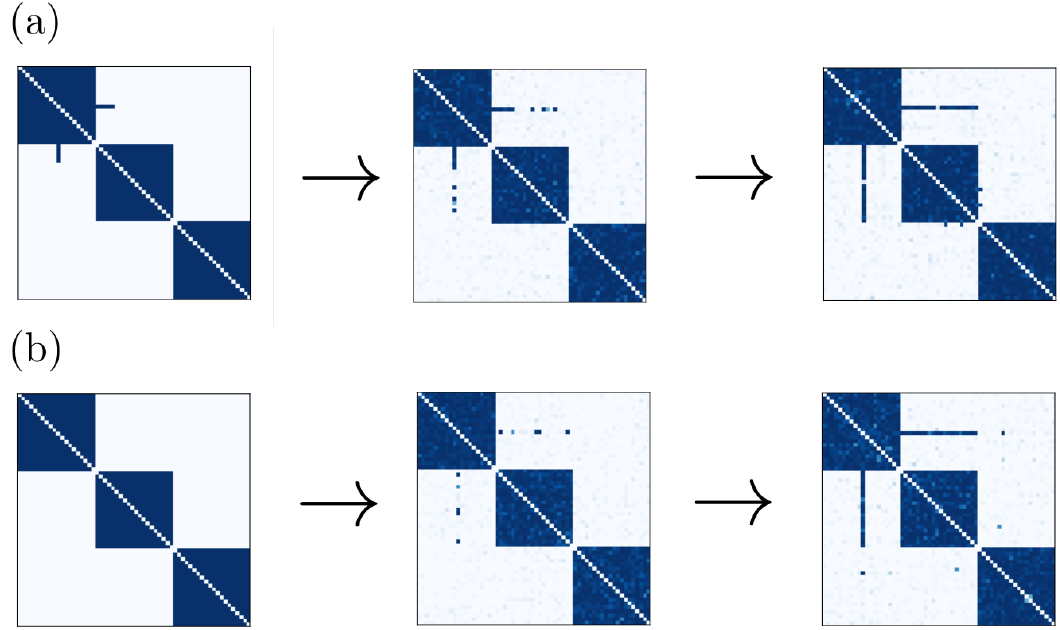
Creation of overlaps. (a): A partial overlap self-completes over time due to plasticity. (b): An overlap emerges for a neuron with lower spontaneous firing rate.

We note that we use a simple analytical rule to determine the spontaneous rates of candidate neurons for the formation of overlaps. To obtain it, we consider two fully connected assemblies of size *N*. We assume that the total firing rate of an overlap neuron, once it is connected to both assemblies, equals the firing rates of the typical neurons, which are only part of a single assembly. If there is an overlap, the firing rates in one assembly will, via the overlap neuron, have a small influence on firing rates in the other assembly. If we neglect this influence, typical neurons’ rates are given by Eq. 14, 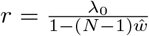. The excess rate 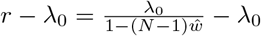 beyond the spontaneous rate *λ*_0_ is thereby due to input from the other neurons in the assembly. An overlap neuron *x* receives twice the input of a typical neuron, because it is part of two assemblies.

Therefore its excess beyond its spontaneous rate is 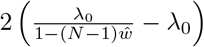. If we call its baseline rate *λ*_*x*,0_, we obtain for the total rate of an overlap neuron

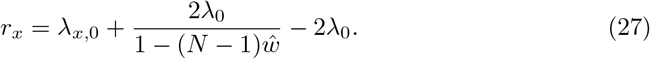

Requiring 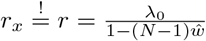 results in

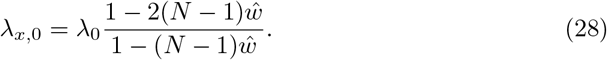

We find that with this spontaneous rate, spontaneous overlap formation is possible and that firing rates for overlap neurons are in good agreement with our predictions, see Fig. S5. The exact threshold for possible spontaneous assembly formation is, however, also dependent on appropriate learning rates and the randomness in the network.

A second way of sustaining overlaps is by having most or all neurons be part of more than one assembly. In this case fusion of two assemblies appears less likely, because all assemblies and neurons are similarly saturated. Indeed, in the brain, we expect most or all neurons to be part of more than one assembly [21, 45]. To test whether our networks are capable of sustaining similarly prominent overlap structures, we consider a network of intertwined assemblies in which each neuron is part of exactly two assemblies. We choose equally sized assemblies of size *n*_A_ and the overlap structure such that for any given assembly each of its neurons is shared with a different assembly. In other words, the overlap between any two assemblies consists of one neuron or a fraction of 1*/n*_A_ of each assembly. This implies that there are in total *n*_A_ + 1 assemblies and that the setup requires a network with *N* = *n*_A_(*n*_A_ + 1)*/*2 neurons. Notably, a network with the same number and size of assemblies, where each neuron is only part of one assembly (no overlap), would require twice as many, (*n*_A_ + 1)*n*_A_, neurons. Fig. 7 shows that this structure is stable under the STDP rule over long timescales. The assemblies have size *n*_A_ = 19; the typical (see the numerical results below) overlap between two assemblies is thus 1*/n*_A_ = 1*/*19 ≈ 5.3%. Random correlations cause additional, extra-assembly connections to appear (more precisely: to strengthen) and intra-assembly connections to disappear (to weaken). To demonstrate that these deviations of individual weights are transient, we have tracked in Fig. 7b and c the strengthening of extra-assembly weights and the weakening of intra-assembly weights, respectively. The upper panel of Fig. 7b displays the sum of the extra-assembly weights over time, indicating that there is no positive drift, i.e. no overall increase of weights that do not belong to the stored assembly structure over time. The lower panel displays the Pearson correlations of the extra-assembly weights over time with the extra-assembly weights observed at five reference time points. With increasing distance to the reference time point, the correlations decay to chance level showing that no persistent pattern of strengthened extra-assembly weights emerges. Fig. 7c analogously displays the differences between the maximal (optimal) and the actual intra-assembly weights. The upper panel shows that there is no overall increase of this difference, i.e. no overall decay of the assembly structure. The lower panel shows that the patterns of weakened intra-assembly weights are transient.

**Fig 7.**
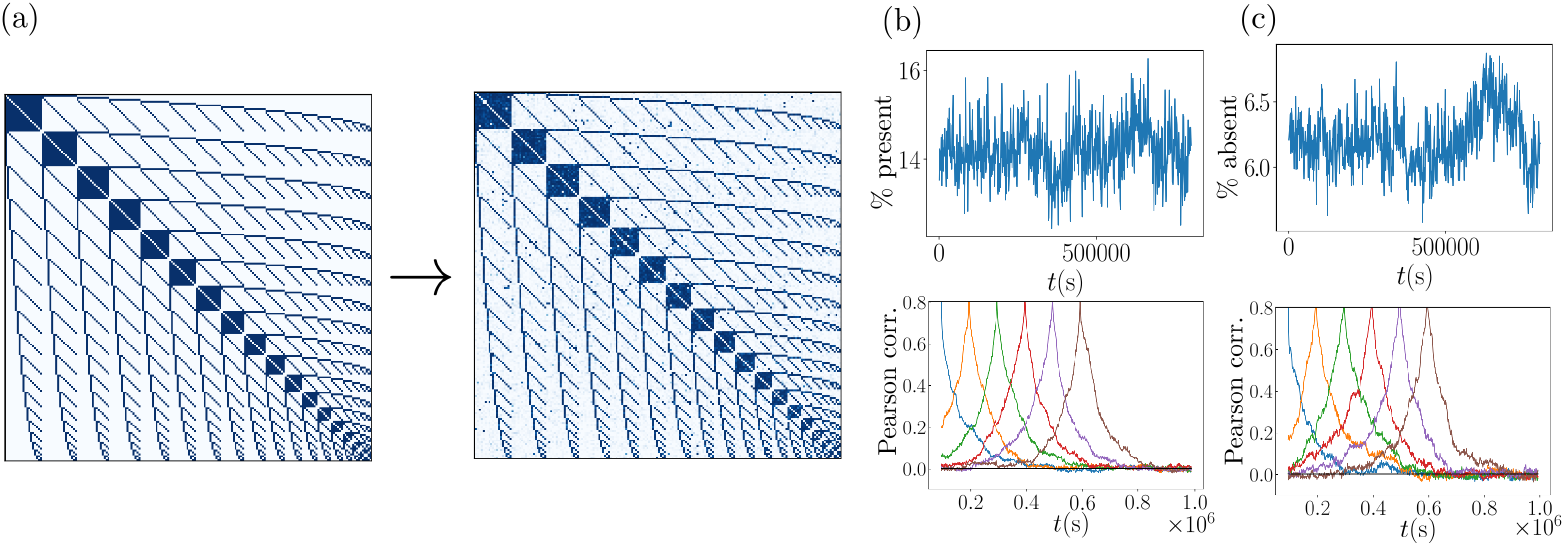
Intertwined assembly network. (a): Initial weight matrix (left hand side) and weight matrix after simulation over 1 × 10^6^ s (right hand side). (b, upper): Sum of the extra-assembly weights, that is of those weights that are zero initially and if the assembly structure is optimally realized. The sum is given as a fraction of the total initial weights. (b, lower): Correlations of the extra-assembly weights with those at six different reference points in time. (c,upper): Sum of the absent intra-assembly weights, that is, the sum of distances of intra-assembly weights from their initial and optimal value *ŵ*. The sum is given as a fraction of the total initial weights. (c,lower): As in (b,lower) for absent intra-assembly weights.

### Drifting assemblies

Experiments have shown that memory representations need not consist of the same neurons over time but can in fact exchange neurons without affecting behavior [25], a phenomenon called representational drift. It may occur because memory assemblies drift, by gradually exchanging neurons between each other [5]. The gradual exchange implies that at each point in time, each assembly is present and unambiguously identifiable by following the course of its evolution from the beginning. In the following, we show that our model networks can give rise to drifting assemblies. The drift happens due to two alternative mechanisms: (i) Neuron exchange between assemblies due to high weight plasticity noise, as in [5], and (ii) formation of temporal overlaps due to modulations in the spontaneous spike rate. Whether drift occurs due to mechanism (i) is chiefly determined by the learning rate *μ*: Fig. 8a displays the assembly dynamics in two networks with different values of *μ*, while all other parameters are kept the same. The network with smaller *μ* has stationary assemblies. In contrast, the network with larger *μ* exhibits drifting assemblies. Specifically, in this network the ensemble of neurons forming an assembly completely changes over time (Fig. S7a). Simultaneously, one can track the identity of an assembly by comparing its constituent neurons over short time intervals; the neurons forming it at one time can be unambiguously matched to the neurons forming it shortly thereafter, since the difference between those two ensembles is still small (Fig. S7b). We can explain the occurrence of this type of drift as an effect of fluctuations in the weight dynamics. The time-averaged weight dynamics as described by 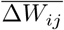 reinforce connections between neurons in the same assembly while suppressing connections between neurons of different assemblies. However, large enough fluctuations in the weight dynamics may nevertheless cause neurons to lose connections within their assembly and form connections to other assemblies. The size of these fluctuations is governed by *μ*, and if they are sufficiently large there is a finite probability that a neuron switches assemblies, see [5] for a detailed discussion. If *μ* is too large, the strong fluctuations prevent assemblies from forming at all.

**Fig 8.**
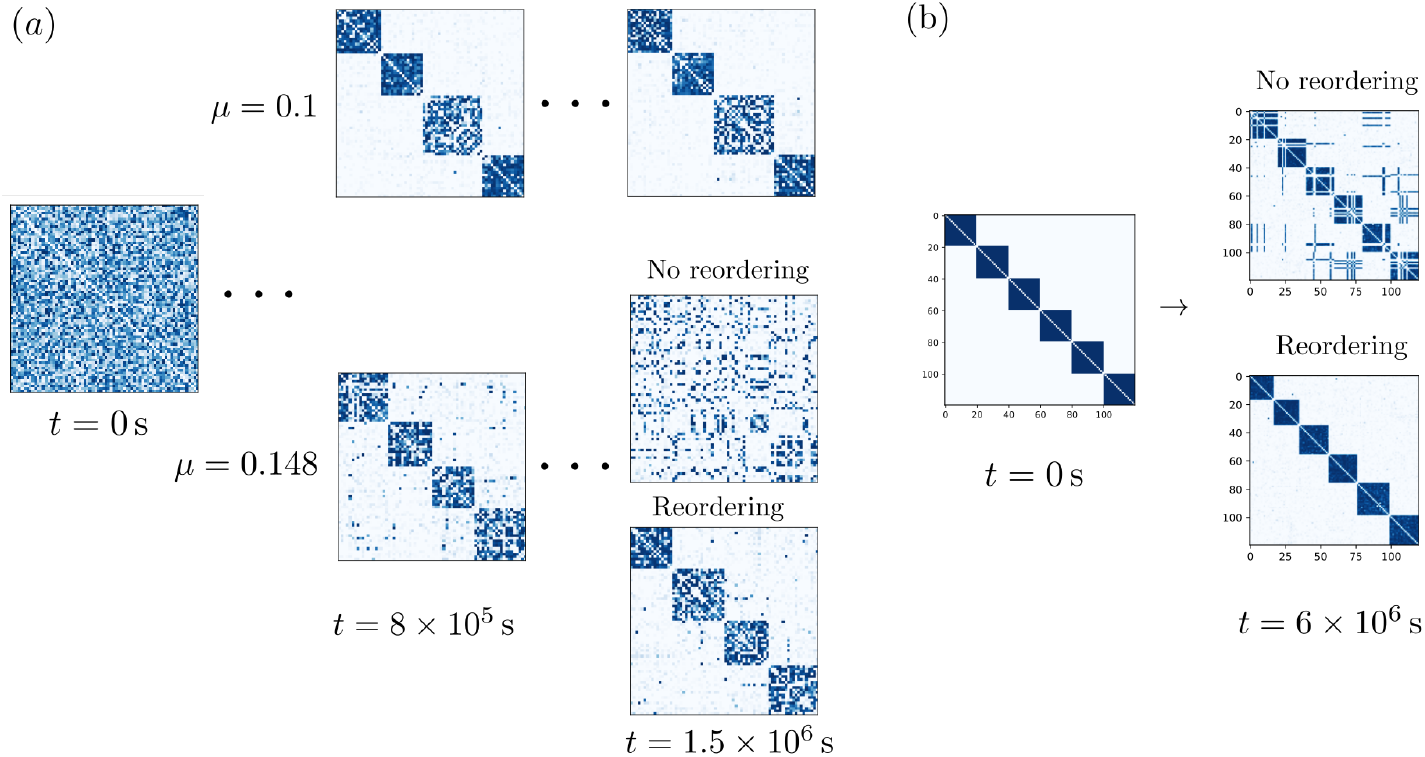
Drifting assemblies. (a): Drift through noisy spiking activity and resulting weight fluctuations. Assembly dynamics for lower (*μ* = 0.1, upper part) and higher (*μ* = 0.145, lower part) learning rate. The panel shows connection matrices *W* at different points in time. Initially, the weight matrices are random (left). After *t* = 8 × 10^5^ s assemblies have spontaneously emerged (middle). The neuron indices are sorted to reveal them (cf. Fig. 2). The simulation is thereafter continued, which shows that for lower learning rate the assemblies are static. In contrast, for higher learning rate the assemblies drift. Since the assemblies exchange neurons, the coupling matrix appears increasingly unstructured. Reordering of the indices, however, reveals that the assemblies are maintained at each point in time. (b): Drift through transient changes in intrinsic neuron properties and resulting transient overlaps. The spontaneous firing rates of neurons change on a slow timescale. This leads to the transient appearance of overlaps, cf. Fig. 6b. When an overlap vanishes, the neuron randomly decides for one of the assemblies, which leads to drifting assemblies.

In the second mechanism switching of assemblies by neurons is a two-step process. In the first step a neuron with a sufficiently low *λ*_0_ spontaneously forms a connection to an additional assembly, as in Fig. 6b. Then, when *λ*_0_ increases again this neuron loses its connections to one of the assemblies. The neuron can thereby leave either of the assemblies it is connected to – if it loses its connections to the assembly that it was originally connected to, it has switched assemblies. If this happens sufficiently often for many neurons in the network, this also causes overall drift on a slow timescale, as shown in Fig. 8b.

### Changing synaptic connectivity

We finally apply our assembly model to networks that undergo changes in their synaptic connectivity. This is motivated by processes related to aging: Anatomical studies have shown that the aging cortex is characterized by a decrease in the number of synaptic spines [26] and presynaptic terminals [27]. These changes may be interpreted as an increase in sparsity of connections between neurons and/or as an overall weakening of synaptic strengths (since connections between neurons often consist of multiple synapses). We can model the former by permanently setting a fraction of entries of the weight matrix, chosen at random, to zero. The latter can be modeled by decreasing *ŵ*. Fig. 3a shows the effect of decreasing *ŵ*: it shifts the zeros of 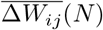 and thus the typical assembly size towards larger *N*. Increased sparsity similarly lowers the branching parameter for an assembly of a given size and thus also shifts the zeros of 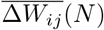 and the characteristic assembly size towards larger *N*, see Fig. 9a. As a result, spontaneously forming assemblies are larger in networks with higher sparsity, see Fig. 9c. In addition Fig. 9c suggests that smaller learning rates further increase the tendency to form larger assemblies. We observe that if the assembly sizes in a network are significantly smaller than the characteristic size predicted by Figs. 3a and 9a, assemblies will merge to form larger ones, see Fig. 9b. This represents a loss in memory capacity. Assuming that in the brain assemblies representing closely related memories merge (due to existing overlaps), during this process the overall memory content becomes less detailed and differentiated. At the same time the neuronal activity during recall increases due to larger assembly sizes indicating a less efficient use of neural resources.

**Fig 9.**
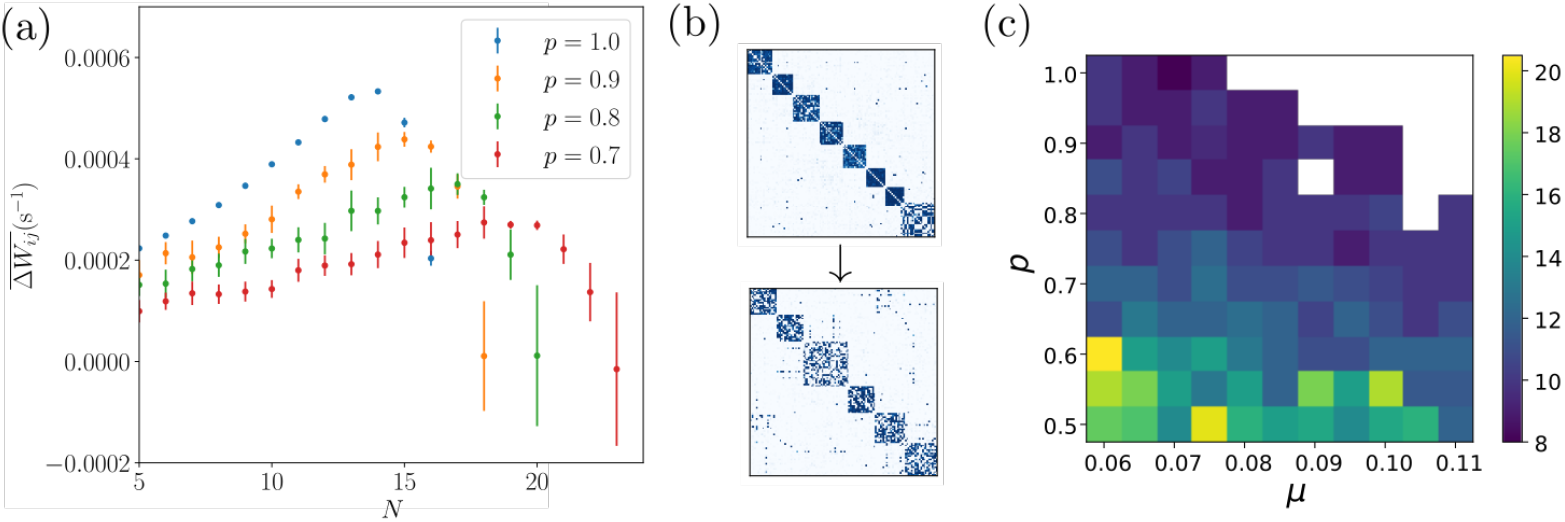
Effects of increasing network sparsity. (a): Time-averaged weight change like in Fig. 3a, of an assembly with intra-assembly connection probability *p*, as a function of its size for different values of *p*. Lower values of *p* lead to a shift of the zero crossing of 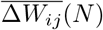 to larger *N*. Simulations with low 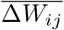 converge slowly, which is reflected by large error bars. (b): An assembly structure becomes coarser (fewer, larger assemblies) as the connectivity sparsity increases. The panel shows the weight matrices of a network in which the probability *p* that a connection between neurons exists is decreased from *p* = 1.0 (upper subpanel, after spontaneous assembly emergence) to *p* = 0.7 (lower subpanel, after re-equilibration). (c): Dependence of the size of a typical assembly on the network connection probability *p* and the learning rate *μ*. The panel shows the median size of spontaneously forming assemblies in networks with random initial connectivity. White areas denote parameter regions for which activity becomes pathological, that is, firing rates diverge.

## Discussion

We have studied assemblies in networks with plasticity that is purely spike-timing dependent. We find that the networks stably store assemblies, which may spontaneously emerge or be learned. Further, the assemblies can have overlaps, spontaneously drift and adapt to changing network properties.

In biological spiking neural networks the change of synaptic efficacies depends to a large extent on STDP, where depression and potentiation are a function of the time lags between pre- and postsynaptic spikes [46–48]. Assemblies may be kept up by the co-spiking of their member neurons, which strengthens their interconnections. If the total input to a neuron is not constrained, large assemblies generate large inputs to each member neuron and thereby highly reliably co-activate them. Furthermore, larger assemblies generate stronger input to other neurons and thereby tend to recruit them, such that these assemblies grow. This generates a positive feedback loop, which can without restraining mechanisms lead to excessive assembly growth. Previous work on assembly networks therefore usually added additional homeostatic plasticity that limits the total input strength to a neuron by fast normalization [5, 7–9, 11–13]. This curtails the resulting excessive assembly growth. The homeostatic plasticity observed in neurobiological experiments is, however, much slower than that required to prevent runaway assembly growth [14–18].

In the present work, we therefore studied assemblies in networks with purely STDP-based plasticity. We find that our depression-dominated STDP-rule (the integral of the STDP window is negative) restricts the growth of assemblies by two mechanisms: On the one hand, the spike rate of the neurons in an assembly grows with the assembly size, which increases the depressing effect of the rate-based term of the time-averaged weight change. On the other hand, the contributions of the higher-order connectivity motifs to the time-averaged weight change become more significant the larger the assemblies are, because longer cascades of spikes become more likely. Further for sufficiently high order the appropriately summed contributions of the motifs are approximately proportional to the integral of the STDP window and thus negative. The contributions of the purely firing rate-based term and the higher-order motifs therefore reduce or even invert the tendency of weights to grow in large assemblies. As a consequence, neurons are more prone to leave a larger assembly and for example join another, smaller one, which induces larger growth of the interconnecting weights. A mechanism similar to the purely firing rate-based effect that we have described has been shown to stabilize the output spike rate of a single neuron receiving feedforward input, since a strong dominance of the rate-based term ultimately leads to the reversal of the sign of plasticity [49]. Another related work, [10], demonstrated the learning of static, non-overlapping assemblies in networks with STDP in the recurrent excitatory and the inhibitory-to-excitatory connections. These assemblies are maintained at least over several hours. The excitatory STDP is thereby balanced (the integral over the STDP window is about zero) such that the inhibitory STDP is effectively much faster. This yields rate homeostasis in the excitatory neurons. In our models the integral over the excitatory STDP curve is negative. Like rate homeostasis, this restricts the maximal average excitatory spike rates. In contrast to rate homeostasis, it also allows for smaller weights, for example in our networks with assembly and background neurons. We note that the fast weight homeostasis in [9] is implemented such that it also only constrains the maximal summed input and output weight of a neuron. We further note that highly diverse average spike rates are observed in biology [50].

We have investigated networks with symmetric plasticity function, motivated by past theoretical work [5, 9] and experimental findings [39, 40]. Experiments have, however, also found strongly asymmetric STDP windows and a dependence of plasticity on details of the neural activity beyond the temporal difference between pairs of spikes [47, 51, 52]. Recent theoretical studies showed assembly formation with such STDP rules and homeostasis [10, 12, 20]. Since the contributions of the higher order network motifs converge generally, for any STDP function, towards a multiple of the integral over the STDP function (cf. Eq. 25), we expect that the limitation of assemblies by our mechanism should also work rather generally. Prerequisites are irregular activity and an STDP function that allows for the growth of small assemblies and has negative integral. It will be important to see whether our mechanisms of assembly growth restriction indeed applies also to networks with strongly asymmetric and more biologically detailed STDP rules.

Our model networks generate irregular, probabilistic spiking activity. This is in agreement with experimentally observed irregular spiking in the cortex. In our models the irregularity of spiking is guaranteed by the usage of a Poisson spiking neuron model. In biological neural networks, it is usually assumed to result from the input fluctuation-driven spiking dynamics of individual neurons, which occur if there is an overall balance of excitatory and inhibitory input. If the excitatory synaptic weights are plastic, as in our model, this balance might be maintained by inhibitory plasticity [6–8, 10]. In the future, it will be important to investigate our mechanisms of assembly growth restriction in networks with explicitly modeled excitatory and inhibitory populations of neurons. Individual neurons then should not intrinsically generate Poisson spiking, but for example be of the integrate-and-fire type.

We observe that our networks are able to maintain prominent overlap structures, where each neuron belongs to more than one assembly. The assembly structure is saturated and remains stable. In particular, additionally increased connections are sparse and transient. Previous related works have not shown similarly prominent overlap structures [5, 7–10, 12, 24], perhaps because the mostly assumed fast homeostatic normalization induces a stronger competition between the assemblies. This may force neurons to decide for one assembly. Networks with overlaps allow a more economic use of neural resources, in the sense that more assemblies of a specific size can be stored, in our example network twice as many as without overlaps. The overlap between assemblies in our simulations is about 5%. This agrees with the overlap estimated for assemblies representing associated concepts in the brain [45]. The overlap of randomly chosen assemblies is smaller, about 1%. Our findings raise a variety of future research questions. We have demonstrated highly prominent overlap structures with symmetric assembly patterns, where any two assemblies have an overlap and all assemblies as well as all overlaps have the same size. Numerical simulations indicate that we can still store similar patterns as these with a few overlaps missing. Future research should address whether and how assembly patterns may be stored that have a similar asymmetry and overlap pattern as estimated for the brain. Inhomogeneous neuron properties such as different spontaneous spike rates may help to achieve this. In this context, it will also be important to investigate how new assemblies can be stored with the right overlaps into a pattern of old ones. Finally, it is unclear whether symmetric or asymmetric assembly patterns with highly prominent overlaps may spontaneously emerge in neural networks. Narrowing down the focus, it will be interesting to study how the overlap between two assemblies evolves and under which conditions it becomes stable. A theoretical description of this evolution needs to account for the back-and-forth interaction between assemblies with dynamically changing overlap. Ref. [45] has recently studied interactions between fixed overlapping assemblies in non-plastic networks. On the microscopic level, our results suggest to investigate in more detail when neurons connect to two assemblies. To address this, one might analytically compute the probabilities of connection weights of a test neuron with two assemblies similar to the description of neuron switching between assemblies in [5]. In contrast to the effectively one-dimensional problem addressed in [5], the absence of homeostatic weight normalization will yield a two-dimensional reduced weight space for our model.

Our model networks can stably maintain assemblies in front of a background of neurons that are not part of any assembly. Such a scenario might be particularly relevant for early development when not many memories have been stored yet. Previous related works usually assume that the entire space is tiled by assemblies [5, 7–10, 12]. The reason may be similar as for the prominent overlap structures, namely that fast homeostatic plasticity has a strong tendency to force neurons into assemblies. Ref. [24] showed assemblies in front of a background of weakly connected neurons in networks with structural plasticity and multiple synapses per connection between neurons.

The assemblies in our networks can drift. Such assembly drift may explain the experimentally observed drift of memory representations in the brain [5, 25]. In our models, assemblies drift by exchanging neurons. In the first model this exchange originates from sufficiently large synaptic weight fluctuations due to noisy spiking, as in [5]. Besides STDP, ref. [5] had to assume the presence of fast weight homeostasis, which helps to keep the representational structure intact. We note that assemblies exchange neurons also in a recent model for spontaneous assembly formation in zebrafish [13], which assumes fast homeostasis as well. The assemblies, however, break down and merge, and are therefore not drifting. In contrast, both our models for drifting assemblies require only STDP as plasticity mechanism. Our second model furthermore introduces a novel neuron exchange and thus drift mechanism: For high spontaneous rate, each neuron belongs to one assembly. If the spontaneous rate of a neuron in the network then transiently drops, the synaptic weights with another assembly increase, such that the neuron belongs to two assemblies. When the intrinsic rate recovers, the synaptic weights with one of the assemblies weaken. This may eliminate the strong weights with the assembly that the neuron originally belonged to and thus induce a switch to the other assembly. The observation suggests a new general mechanism for representational drift in the brain: Transient (or persistent) changes in single neuron properties may lead to changes in the synaptic weights. These in turn induce a change in the features represented by the neuron. The changes in the synaptic weights and the representation may thereby be much longer lasting than the changes in the intrinsic neuronal properties.

Since in the aging brain the overall connectivity decreases [26, 27], we have explored the impact of a such a decrease on the assemblies in our model networks. We observe that the size of assemblies is inversely related to the connection probability in the network. We expect that a similar relation can be observed rather generally, for example also in model networks where the assemblies are stabilized by fast homeostasis. In particular, sparser networks should, all other parameters being equal, lead to larger assemblies that are activated during recall. This is consistent with the observation that neural activity for the same task is stronger in aged individuals [53], where the neural networks are more sparsely connected. An increased assembly size and the resulting stronger activity during reactivation might also explain why episodic memories are experienced more vividly in elderly subjects [54]. In our model networks, we observe that assemblies merge to larger ones if networks become increasingly sparse. Such mergers might explain why episodic memories become less detailed in the aging brain [54, 55].

## Supporting information

S1 Appendix: Supporting analysis and figures

## Acknowledgments

We thank Sven Goedeke for helpful comments on the manuscript.

## Notes

### Competing Interest Statement

The authors have declared no competing interest.

### Summary of Updates

Main text: additions to section 'Plasticity in homogeneously connected assemblies' and 'Storing new assemblies', added Figure 3b, further clarifications throughout the text Supplement: further detail to text S2, added Figures S2, S3, S4, S5,

