## Supplementary material for "Purely STDP-based assembly dynamics: stability, learning, overlaps, drift and aging": S1 Appendix: Supporting analysis and figures

### S.1 Time-averaged weight change in fully connected assemblies

The correlation function of our Poisson model neurons reads in frequency space

$$\tilde{C}(\omega) = \tilde{C}^0(\omega) + \tilde{C}^1(\omega) = 2\pi\delta(\omega) r r^T + (\mathbb{1} - \tilde{a}(\omega)W)^{-1} D \left(\mathbb{1} - \tilde{a}(-\omega)W^T\right)^{-1}, \tag{S.1}$$

see main text Eq. 4. For a homogeneously coupled assembly of $N$ neurons with recurrent weights $\hat{w}$ and identical spontaneous rates $\lambda_0$, all rates $r_i$ are the same,

$$r_i =: r = \frac{\lambda_0}{1-(N-1)\hat{w}}, \tag{S.2}$$

see Eq. 14. This fixes the first right hand side term of Eq. S.1 and $D = r\mathbb{1}$. $W$ has zeros on the diagonal and otherwise entries $\hat{w}$. The second right hand side term of Eq. S.1 can thus be written as

$$\begin{aligned}\tilde{C}^1(\omega) &= (\mathbb{1} - \tilde{a}(\omega)W)^{-1} D \left(\mathbb{1} - \tilde{a}(-\omega)W^T\right)^{-1} \\ &= r\left(\alpha_+\mathbb{1} + \beta_+ J_N\right)^{-1}\left(\alpha_-\mathbb{1} - \beta_+ J_N\right)^{-1},\end{aligned} \tag{S.3}$$

where $J_N$ is the $N \times N$ matrix of ones, $\alpha_\pm = 1 + \hat{w}\tilde{a}(\pm\omega)$, and $\beta_\pm = -\hat{w}\tilde{a}(\pm\omega)$. We obtain the inverses of the matrices by assuming that they have the same general structure, that is using the ansatz

$$(\alpha_\pm\mathbb{1} + \beta_\pm J_N)^{-1} = \gamma_\pm\mathbb{1} + \delta_\pm J_N. \tag{S.4}$$

This yields

$$\gamma_\pm = \frac{1}{\alpha_\pm} = \frac{1}{1+\hat{w}\tilde{a}(\pm\omega)}, \tag{S.5}$$

$$\delta_\pm = \frac{-\beta_\pm}{\alpha_\pm(\alpha_\pm + N\beta_\pm)} = \frac{\hat{w}\tilde{a}(\pm\omega)}{(1+\hat{w}\tilde{a}(\pm\omega))(1-\hat{w}\tilde{a}(\pm\omega)(N-1))}. \tag{S.6}$$

In terms of $\gamma_\pm$ and $\delta_\pm$, $C^1(\omega)$ reads

$$\begin{aligned}\tilde{C}^1(\omega) &= r\left(\gamma_+\mathbb{1} + \delta_+ J_N\right)\left(\gamma_-\mathbb{1} + \delta_- J_N\right) \\ &= r\left(\gamma_+\gamma_-\mathbb{1} + \gamma_+\delta_- J_N + \gamma_-\delta_+ J_N + N\delta_+\delta_- J_N\right).\end{aligned} \tag{S.7}$$

Since there are no autapses in the network, we are only interested in the off-diagonal elements,

$$\tilde{C}^1_{i\neq j}(\omega) = r\left(\gamma_+\delta_- + \gamma_-\delta_+ + N\delta_+\delta_-\right). \tag{S.8}$$

The Fourier transform of the synaptic kernel Eq. 2 is

$$\tilde{a}(\omega) = \frac{1}{1+i\omega\tau_\mathrm{s}}, \tag{S.9}$$

such that

$$\begin{aligned}\tilde{C}^1_{i\neq j}(\omega) = r\Bigg(&\frac{\hat{w}(1+i\omega\tau_\mathrm{s})(1-i\omega\tau_\mathrm{s})}{(1+i\omega\tau_\mathrm{s}+\hat{w})(1-i\omega\tau_\mathrm{s}+\hat{w})(1-i\omega\tau_\mathrm{s}-(N-1)\hat{w})} + \\ &+ \frac{\hat{w}(1-i\omega\tau_\mathrm{s})(1+i\omega\tau_\mathrm{s})}{(1-i\omega\tau_\mathrm{s}+\hat{w})(1+i\omega\tau_\mathrm{s}+\hat{w})(1+i\omega\tau_\mathrm{s}-(N-1)\hat{w})} + \\ &+ \frac{N\hat{w}^2(1+i\omega\tau_\mathrm{s})(1-i\omega\tau_\mathrm{s})}{(1+i\omega\tau_\mathrm{s}+\hat{w})(1+i\omega\tau_\mathrm{s}-(N-1)\hat{w})(1-i\omega\tau_\mathrm{s}+\hat{w})(1-i\omega\tau_\mathrm{s}-(N-1)\hat{w})}\Bigg).\end{aligned} \tag{S.10}$$

For the Fourier transform of the plasticity window Eq. 5 we have

$$\tilde{F}(\omega) = \frac{2A_\mathrm{p}\tau_\mathrm{p}}{(1+i\tau_\mathrm{p}\omega)(1-i\tau_\mathrm{p}\omega)} + \frac{2A_\mathrm{d}\tau_\mathrm{d}}{(1+i\tau_\mathrm{d}\omega)(1-i\tau_\mathrm{d}\omega)} = \tilde{F}(-\omega). \tag{S.11}$$

Inserting Eqs. S.1 and S.2 into Eq. 8 gives

$$\begin{aligned} \overline{\Delta W_{ij}}(N) &= \frac{1}{2\pi}\int_{-\infty}^{\infty} \mathrm{d}\omega\ \tilde{C}_{ij}(\omega)\tilde{F}(-\omega) \\ &= \frac{f_0\lambda_0^2}{(1-(N-1)\hat{w})^2} + \frac{1}{2\pi}\int_{-\infty}^{\infty}\mathrm{d}\omega\ \tilde{C}^1_{ij}(\omega)\tilde{F}(-\omega). \end{aligned} \tag{S.12}$$

Inserting Eqs. S.2, S.10 and S.11 into Eq. S.12 results in

$$\begin{aligned} \frac{1}{2\pi}\int_{-\infty}^{\infty}\mathrm{d}\omega\ \tilde{C}^1_{ij}(\omega)\tilde{F}(-\omega) = \frac{1}{2\pi}\int_{-\infty}^{\infty}\mathrm{d}\omega\ &\left(2\pi\frac{\hat{w}(1+i\omega\tau_\mathrm{s})(1-i\omega\tau_\mathrm{s})}{(1+i\omega\tau_\mathrm{s}+\hat{w})(1-i\omega\tau_\mathrm{s}+\hat{w})(1-i\omega\tau_\mathrm{s}-(N-1)\hat{w})}+\right. \\ &+\frac{\hat{w}(1-i\omega\tau_\mathrm{s})(1+i\omega\tau_\mathrm{s})}{(1-i\omega\tau_\mathrm{s}+\hat{w})(1+i\omega\tau_\mathrm{s}+\hat{w})(1+i\omega\tau_\mathrm{s}-(N-1)\hat{w})}+ \\ &\left.+\frac{N\hat{w}^2(1+i\omega\tau_\mathrm{s})(1-i\omega\tau_\mathrm{s})}{(1+i\omega\tau_\mathrm{s}+\hat{w})(1+i\omega\tau_\mathrm{s}-(N-1)\hat{w})(1-i\omega\tau_\mathrm{s}+\hat{w})(1-i\omega\tau_\mathrm{s}-(N-1)\hat{w})}\right)\times \\ &\times\left(\frac{2A_\mathrm{p}\tau_\mathrm{p}}{(1+i\tau_\mathrm{p}\omega)(1-i\tau_\mathrm{p}\omega)}+\frac{2A_\mathrm{d}\tau_\mathrm{d}}{(1+i\tau_\mathrm{d}\omega)(1-i\tau_\mathrm{d}\omega)}\right)\frac{\lambda_0}{1-(N-1)\hat{w}}. \end{aligned} \tag{S.13}$$

This integral can be straightforwardly computed using the residue theorem. Together with the zeroth-order term given by Eq. S.12 we obtain the following closed-form expression for the time-averaged weight change:

$$\begin{aligned} \overline{\Delta W_{ij}}(N) = &\frac{2\lambda_0^2(A_\mathrm{p}\tau_\mathrm{p}+A_\mathrm{d}\tau_\mathrm{d})}{(1-(N-1)\hat{w})^2}+ \\ &+\frac{\lambda_0 A_\mathrm{p}\tau_\mathrm{p}\hat{w}\left[(2-(N-2)\hat{w})\tau_\mathrm{p}+(2-(N-2)\hat{w}-(N-1)\hat{w}^2)\tau_\mathrm{s}\right]}{(1+\hat{w})(1-(N-1)\hat{w})^2(\tau_\mathrm{s}+(1+\hat{w})\tau_\mathrm{p})(\tau_\mathrm{s}+(1-(N-1)\hat{w})\tau_\mathrm{p})}+ \\ &+\frac{\lambda_0 A_\mathrm{d}\tau_\mathrm{d}\hat{w}\left[(2-(N-2)\hat{w})\tau_\mathrm{d}+(2-(N-2)\hat{w}-(N-1)\hat{w}^2)\tau_\mathrm{s}\right]}{(1+\hat{w})(1-(N-1)\hat{w})^2(\tau_\mathrm{s}+(1+\hat{w})\tau_\mathrm{d})(\tau_\mathrm{s}+(1-(N-1)\hat{w})\tau_\mathrm{d})}, \end{aligned} \tag{S.14}$$

main text Eq. 15. The first right hand side term term is negative for all admissible $N$ because for our networks $f_0 < 0$. Due to the restriction $N < 1+\frac{1}{\hat{w}}$ the terms $2-(N-2)\hat{w}$, $2-(N-2)\hat{w}-(N-1)\hat{w}^2$, and $(1-(N-1)\hat{w})$, which occur in the numerators and denominators of the second and third right hand side terms, are positive. Therefore the second right hand side term is positive, while the third one is negative since $A_\mathrm{d} < 0$.

To show that for $\hat{w} \to 0$ we have $\overline{\Delta W_{ij}}(N) < 0$ for all $N$, we first isolate the always positive factor $1/(1-(N-1)\hat{w})^2$ in Eq. S.14,

$$\begin{aligned} \overline{\Delta W_{ij}}(N) = &\left(2\lambda_0^2(A_\mathrm{p}\tau_\mathrm{p}+A_\mathrm{d}\tau_\mathrm{d})+\right. \\ &+\frac{\lambda_0 A_\mathrm{p}\tau_\mathrm{p}\hat{w}\left[(2-(N-2)\hat{w})\tau_\mathrm{p}+(2-(N-2)\hat{w}-(N-1)\hat{w}^2)\tau_\mathrm{s}\right]}{(1+\hat{w})(\tau_\mathrm{s}+(1+\hat{w})\tau_\mathrm{p})(\tau_\mathrm{s}+(1-(N-1)\hat{w})\tau_\mathrm{p})}+ \\ &\left.+\frac{\lambda_0 A_\mathrm{d}\tau_\mathrm{d}\hat{w}\left[(2-(N-2)\hat{w})\tau_\mathrm{d}+(2-(N-2)\hat{w}-(N-1)\hat{w}^2)\tau_\mathrm{s}\right]}{(1+\hat{w})(\tau_\mathrm{s}+(1+\hat{w})\tau_\mathrm{d})(\tau_\mathrm{s}+(1-(N-1)\hat{w})\tau_\mathrm{d})}\right) \\ &\times\frac{1}{(1-(N-1)\hat{w})^2}. \end{aligned} \tag{S.15}$$

The terms within the large bracket are finite for all admissible $N$. The first summand in the large bracket is negative and independent of $N$ and $\hat{w}$. To obtain an upper bound for the large bracket, we omit the negative third summand. Further, we increase the numerator of the second summand by inserting the lower bound $N = 0$ for $N$. We decrease the denominator by inserting the upper bound $N = 1 + \frac{1}{\hat{w}}$. This results in

$$\begin{aligned}\overline{\Delta W_{ij}}(N) < \Bigg( & 2\lambda_0^2(A_\mathrm{p}\tau_\mathrm{p} + A_\mathrm{d}\tau_\mathrm{d}) + \\ & + \frac{\lambda_0 A_\mathrm{p}\tau_\mathrm{p}\hat{w}\left[(2+2\hat{w})\tau_\mathrm{p} + (2+2\hat{w}+\hat{w}^2)\tau_\mathrm{s}\right]}{(1+\hat{w})(\tau_\mathrm{s} + (1+\hat{w})\tau_\mathrm{p})\tau_\mathrm{s}}\Bigg) \\ & \times \frac{1}{(1-(N-1)\hat{w})^2}. \end{aligned} \tag{S.16}$$

Due to the prefactor $\hat{w}$, the positive second summand will go to zero with $\hat{w}$. In particular its magnitude will at some point fall below the magnitude of the negative first summand, such that $\overline{\Delta W_{ij}}(N) < 0$ for all $N$, see also Fig. S3.

### S.2 Asymptotic behavior of motif contributions to time-averaged plasticity

We first show that the series expansion Eq. 11 of the time-averaged weight change in a homogeneously connected assembly simplifies to

$$\begin{aligned}\overline{\Delta W_{ij}} &= f_0 r_i r_j + \sum_{\alpha,\beta} f_{\alpha\beta} \sum_m r_m (W^\alpha)_{im} (W^\beta)_{jm} \\ &= f_0 r^2 + \frac{r}{N} \sum_{k=1}^{\infty} f_k \left((N-1)^k - (-1)^k\right) \hat{w}^k, \end{aligned} \tag{S.17}$$

where

$$f_k := \sum_{\alpha+\beta=k} f_{\alpha\beta}. \tag{S.18}$$

In homogeneous assemblies we have $r_i = r$ and $W = \hat{w}(J_N - \mathbb{1})$. Eq. 11 thus yields

$$\begin{aligned}\overline{\Delta W_{ij}} &= f_0 r^2 + r \sum_{\alpha,\beta} f_{\alpha\beta} \left(J_N - \mathbb{1}\right)_{ij}^{\alpha+\beta} \hat{w}^{\alpha+\beta} \\ &= f_0 r^2 + r \sum_{k=1}^{\infty} f_k \left(J_N - \mathbb{1}\right)_{ij}^{k} \hat{w}^{k}. \end{aligned} \tag{S.19}$$

We observe that $J_N^m = N^{m-1} J_N$ for $m \geq 1$ while $J_N^0 = \mathbb{1}$. With this the binomial in Eq. S.19 can be expanded to

$$\begin{aligned}\left(J_N - \mathbb{1}\right)^k &= \sum_{l=0}^{k} \binom{k}{l} J_N^l \left(-\mathbb{1}\right)^{k-l} \\ &= \sum_{l=0}^{k} \binom{k}{l} N^{l-1} J_N (-1)^{k-l} - \frac{(-1)^k}{N} J_N + \left(-\mathbb{1}\right)^k. \end{aligned} \tag{S.20}$$

We are again interested only in off-diagonal elements:

$$\begin{aligned}\left(J_N - \mathbb{1}\right)_{i\neq j}^k &= \sum_{l=0}^{k} \binom{k}{l} N^{l-1} (-1)^{k-l} - \frac{(-1)^k}{N} \\ &= \frac{(N-1)^k - (-1)^k}{N}. \end{aligned} \tag{S.21}$$

Inserting this into Eq. S.19 gives Eq. S.17.

We now show that

$$\lim_{k\to\infty}\sum_{\alpha+\beta=k} f_{\alpha\beta} = \frac{f_0}{2\tau_\mathrm{s}}. \tag{S.22}$$

We start by using Eq. 12, to write

$$\sum_{\alpha+\beta=k} f_{\alpha\beta} = \frac{1}{2\pi}\int_{-\infty}^{\infty} \mathrm{d}\omega\ \tilde{F}(-\omega) \sum_{\alpha+\beta=k} \tilde{a}(\omega)^\alpha \tilde{a}(-\omega)^\beta. \tag{S.23}$$

The sum in the integrand can be rewritten using Eq. S.9,

$$\begin{aligned}
\sum_{\alpha+\beta=k} \tilde{a}(\omega)^\alpha \tilde{a}(-\omega)^\beta &= \sum_{\alpha=0}^{k} \tilde{a}(\omega)^\alpha \tilde{a}(-\omega)^{k-\alpha} \\
&= \sum_{\alpha=0}^{k} \left(\frac{1}{1+i\omega\tau_\mathrm{s}}\right)^\alpha \left(\frac{1}{1-i\omega\tau_\mathrm{s}}\right)^{k-\alpha} \\
&= \sum_{\alpha=0}^{k} \left(\frac{1-i\omega\tau_\mathrm{s}}{1+i\omega\tau_\mathrm{s}}\right)^\alpha \left(\frac{1}{1-i\omega\tau_\mathrm{s}}\right)^{k} \\
&= \frac{1-\left(\frac{1-i\omega\tau_\mathrm{s}}{1+i\omega\tau_\mathrm{s}}\right)^{k+1}}{1-\left(\frac{1-i\omega\tau_\mathrm{s}}{1+i\omega\tau_\mathrm{s}}\right)} \left(\frac{1}{1-i\omega\tau_\mathrm{s}}\right)^{k} \\
&= \frac{\omega\tau_\mathrm{s}-i}{2\omega\tau_\mathrm{s}}\left(\frac{1}{1-i\omega\tau_\mathrm{s}}\right)^{k} + \frac{\omega\tau_\mathrm{s}+i}{2\omega\tau_\mathrm{s}}\left(\frac{1}{1+i\omega\tau_\mathrm{s}}\right)^{k}.
\end{aligned} \tag{S.24}$$

We substitute $\omega' := \omega\tau_\mathrm{s}$ and insert Eq. S.24 into Eq. S.23:

$$\begin{aligned}
\sum_{\alpha+\beta=k} f_{\alpha\beta} &= \frac{1}{2\pi\tau_\mathrm{s}}\int_{-\infty}^{\infty} \mathrm{d}\omega'\ \tilde{F}(-\omega'/\tau_\mathrm{s}) \left(\frac{\omega'-i}{2\omega'}\left(\frac{1}{1-i\omega'}\right)^{k} + \frac{\omega'+i}{2\omega'}\left(\frac{1}{1+i\omega'}\right)^{k}\right) \\
&= \frac{1}{2\pi\tau_\mathrm{s}}\int_{-\infty}^{\infty} \mathrm{d}\omega' \int_{-\infty}^{\infty} \mathrm{d}t\ \mathrm{e}^{\frac{i\omega' t}{\tau_\mathrm{s}}} F(t) \left(\frac{\omega'-i}{2\omega'}\left(\frac{1}{1-i\omega'}\right)^{k} + \frac{\omega'+i}{2\omega'}\left(\frac{1}{1+i\omega'}\right)^{k}\right) \\
&= \frac{1}{2\pi\tau_\mathrm{s}}\int_{-\infty}^{\infty} \mathrm{d}t\ F(t) \int_{-\infty}^{\infty} \mathrm{d}\omega'\ \mathrm{e}^{\frac{i\omega' t}{\tau_\mathrm{s}}} \left(\frac{\omega'-i}{2\omega'}\left(\frac{1}{1-i\omega'}\right)^{k} + \frac{\omega'+i}{2\omega'}\left(\frac{1}{1+i\omega'}\right)^{k}\right).
\end{aligned} \tag{S.25}$$

In the second line we employed the definition of the Fourier transform to substitute $\tilde{F}(-\omega/\tau_\mathrm{s})$. (Alternatively, one could use Plancherel's theorem and the inverse Fourier transform of S.24 to obtain the third line.) The integrand of the inner integral of Eq. S.25 has poles at $\omega' = \pm i$ (the singularity at $\omega' = 0$ is removable). For $t > 0$ ($t < 0$) we can compute the integral using a contour over the upper (lower) half complex plane. The residue theorem then yields

$$\begin{aligned}
&\frac{1}{2\pi\tau_\mathrm{s}}\int_{-\infty}^{\infty} \mathrm{d}t\ F(t) \int_{-\infty}^{\infty} \mathrm{d}\omega'\ \mathrm{e}^{\frac{i\omega' t}{\tau_\mathrm{s}}} \left(\frac{\omega'-i}{2\omega'}\left(\frac{1}{1-i\omega'}\right)^{k} + \frac{\omega'+i}{2\omega'}\left(\frac{1}{1+i\omega'}\right)^{k}\right) \\
&= -\frac{i}{\tau_\mathrm{s}}\int_{-\infty}^{0} \mathrm{d}t\ F(t)\,\mathrm{Res}\left(\mathrm{e}^{\frac{i\omega' t}{\tau_\mathrm{s}}}\ \frac{\omega'-i}{2\omega'}\left(\frac{1}{1-i\omega'}\right)^{k}, -i\right) + \frac{i}{\tau_\mathrm{s}}\int_{0}^{\infty} \mathrm{d}t\ F(t)\,\mathrm{Res}\left(\mathrm{e}^{\frac{i\omega' t}{\tau_\mathrm{s}}}\ \frac{\omega'+i}{2\omega'}\left(\frac{1}{1+i\omega'}\right)^{k}, i\right)
\end{aligned} \tag{S.26}$$

We use that

$$\mathrm{Res}(g,c) = \frac{1}{(k-1)!}\lim_{z\to c}\frac{\mathrm{d}^{k-1}}{\mathrm{d}z^{k-1}}((z-c)^k g(z)) \tag{S.27}$$

if $g(z)$ has a kth order pole at $z=c$. For $\alpha+\beta=k+1$ the residue at $\omega'=+i$ in Eq. S.26 thus becomes

$$\begin{aligned}
\mathrm{Res}\left(\mathrm{e}^{\frac{i\omega' t}{\tau_\mathrm{s}}}\frac{\omega'+i}{2\omega'}\left(\frac{1}{1+i\omega'}\right)^{k+1}, i\right) &= \frac{1}{k!}\lim_{\omega'\to i}\frac{\mathrm{d}^k}{\mathrm{d}\omega'^k}\left((\omega'-i)^{k+1}\left(\frac{1}{1+i\omega'}\right)^{k+1}\left(\frac{1}{2}+\frac{i}{2\omega'}\right)\mathrm{e}^{\frac{i\omega' t}{\tau_\mathrm{s}}}\right)\\
&= \frac{1}{i^{k+1}k!}\lim_{\omega'\to i}\frac{\mathrm{d}^k}{\mathrm{d}\omega'^k}\left(\left(\frac{1}{2}+\frac{i}{2\omega'}\right)\mathrm{e}^{\frac{i\omega' t}{\tau_\mathrm{s}}}\right)\\
&= \frac{1}{i^{k+1}k!}\lim_{\omega'\to i}\left(\frac{1}{2}\left(\frac{it}{\tau_\mathrm{s}}\right)^k\mathrm{e}^{\frac{i\omega' t}{\tau_\mathrm{s}}}+\sum_{j=0}^{k}\binom{k}{j}\left(\frac{it}{\tau_\mathrm{s}}\right)^j\mathrm{e}^{\frac{i\omega' t}{\tau_\mathrm{s}}}\frac{(-1)^{k-j}(k-j)!\,i}{2\omega'^{k-j+1}}\right)\\
&= \frac{1}{i^{k+1}}\lim_{\omega'\to i}\left(\frac{1}{2k!}\left(\frac{it}{\tau_\mathrm{s}}\right)^k\mathrm{e}^{\frac{i\omega' t}{\tau_\mathrm{s}}}+\sum_{j=0}^{k}\left(\frac{it}{\tau_\mathrm{s}}\right)^j\mathrm{e}^{\frac{i\omega' t}{\tau_\mathrm{s}}}\frac{(-1)^{k-j}\,i}{2\,j!\,\omega'^{k-j+1}}\right)\\
&= \frac{\mathrm{e}^{-\frac{t}{\tau_\mathrm{s}}}}{2ik!}\left(\frac{t}{\tau_\mathrm{s}}\right)^k+\frac{1}{2i}\mathrm{e}^{-\frac{t}{\tau_\mathrm{s}}}\sum_{j=0}^{k}\frac{1}{j!}\left(\frac{t}{\tau_\mathrm{s}}\right)^j.
\end{aligned} \tag{S.28}$$

Similarly for the residue at $\omega'=-i$ we obtain

$$\mathrm{Res}\left(\mathrm{e}^{\frac{i\omega' t}{\tau_\mathrm{s}}}\frac{\omega'-i}{2\omega'}\left(\frac{1}{1-i\omega'}\right)^{k+1}, -i\right) = \frac{i\,\mathrm{e}^{\frac{t}{\tau_\mathrm{s}}}}{2k!}\left(\frac{-t}{\tau_\mathrm{s}}\right)^k+\frac{i}{2}\mathrm{e}^{\frac{t}{\tau_\mathrm{s}}}\sum_{j=0}^{k}\frac{1}{j!}\left(\frac{-t}{\tau_\mathrm{s}}\right)^j. \tag{S.29}$$

Inserting Eq. S.28 and Eq. S.29 into Eq. S.26 gives

$$\sum_{\alpha+\beta=k} f_{\alpha\beta} = \frac{1}{\tau_\mathrm{s}}\int_{-\infty}^{\infty}\mathrm{d}t\;F(t)\left(\frac{\mathrm{e}^{-\frac{|t|}{\tau_\mathrm{s}}}}{2(k-1)!}\left(\frac{|t|}{\tau_\mathrm{s}}\right)^{k-1}+\frac{1}{2}\mathrm{e}^{-\frac{|t|}{\tau_\mathrm{s}}}\sum_{j=0}^{k-1}\frac{1}{j!}\left(\frac{|t|}{\tau_\mathrm{s}}\right)^j\right) \tag{S.30}$$

$$=: \frac{1}{\tau_\mathrm{s}}\int_{-\infty}^{\infty}\mathrm{d}t\;F(t)A_k(t). \tag{S.31}$$

We then take the limit $k\to\infty$:

$$\begin{aligned}
\lim_{k\to\infty}\sum_{\alpha+\beta=k} f_{\alpha\beta} &= \lim_{k\to\infty}\frac{1}{\tau_\mathrm{s}}\int_{-\infty}^{\infty}\mathrm{d}t\;F(t)\left(\frac{\mathrm{e}^{-\frac{|t|}{\tau_\mathrm{s}}}}{2(k-1)!}\left(\frac{|t|}{\tau_\mathrm{s}}\right)^{k-1}+\frac{1}{2}\mathrm{e}^{-\frac{|t|}{\tau_\mathrm{s}}}\sum_{j=0}^{k-1}\frac{1}{j!}\left(\frac{|t|}{\tau_\mathrm{s}}\right)^j\right)\\
&= \lim_{k\to\infty}\frac{1}{\tau_\mathrm{s}}\int_{-\infty}^{\infty}\mathrm{d}t\;F(t)\frac{\mathrm{e}^{-\frac{|t|}{\tau_\mathrm{s}}}}{2(k-1)!}\left(\frac{|t|}{\tau_\mathrm{s}}\right)^{k-1}+\frac{1}{2\tau_\mathrm{s}}\int_{-\infty}^{\infty}\mathrm{d}t\;F(t)\;\mathrm{e}^{-\frac{|t|}{\tau_\mathrm{s}}}\lim_{k\to\infty}\sum_{j=0}^{k-1}\frac{1}{j!}\left(\frac{|t|}{\tau_\mathrm{s}}\right)^j.
\end{aligned} \tag{S.32}$$

The first limit vanishes as long as $F(t)$ goes to zero polynomially or faster for large $|t|$. This is guaranteed if $F(t)$ is integrable over $(-\infty,\infty)$, which we already implicitly assume for example to do the Fourier transform. The limit of the power series in the second term is the exponential function. (Limit and integral are there interchangeable due to the dominated convergence theorem.) From Eq. S.32 we thus obtain the final result:

$$\begin{aligned}\lim_{k\to\infty}\sum_{\alpha+\beta=k} f_{\alpha\beta} &= \frac{1}{2\tau_\mathrm{s}}\int_{-\infty}^{\infty}\mathrm{d}t\ F(t)\ \mathrm{e}^{-\frac{|t|}{\tau_\mathrm{s}}}\mathrm{e}^{\frac{|t|}{\tau_\mathrm{s}}}\\ &= \frac{f_0}{2\tau_\mathrm{s}},\end{aligned} \tag{S.33}$$

see also Fig. S1a.

To develop an intuition about this result, we observe that the inner integral of Eq. S.25 is up to a constant factor the inverse Fourier transformation of $\sum_{\alpha+\beta=k}\tilde{a}(\omega)^\alpha\tilde{a}(-\omega)^\beta$. Using its linearity we may apply the inverse Fourier transform to each summand individually. This yields $a(t)$ convolved $\alpha$ times with itself and with $a(-t)$ convolved $\beta$ times with itself. Due to Eq. 2 this equals the probability distribution of a sum of $\alpha$ exponentially distributed independent stochastic variables with mean $\tau_s$ minus the sum of $\beta$ stochastic variables of the same type. In other words, we have the probability distribution of a spike time occurring at the end of a cascade of $\alpha$ spikes with exponentially distributed inter-spike intervals minus the time of a spike occurring at the end of a similar cascade of $\beta$ spikes. This reflects the fact that the term $f_{\alpha\beta}$ covers the impact of the motif where a neuron evokes a spike in the pre- and postsynaptic neurons after spike cascades of length $\beta$ and $\alpha$. The probability distribution has mean $(\alpha-\beta)\tau_s$ and standard deviation $\sqrt{k}\tau_s$. The standard deviations of all distributions are thus identical and neighboring distributions have distance $2\tau_s$. According to the central limit theorem, for large $k$ the distributions approximate normal distributions. For $\alpha$ and $\beta$ adding to the same $k$, these are equidistantly shifted but otherwise identical, see Fig. S1c. With increasing $k$ they broaden, such that their superposition forms a plateau, see Fig. S1b. The increase in overlap thereby compensates the decrease in the distribution heights. The number of distributions increases with increasing $k$. The added distributions, however, do not lead to a non-compensatory increase of the superposition in the relevant center where it overlaps with $F$. This is because the added distributions initially do not reach the center, as their mean scales with $k$ while their width scales only with $\sqrt{k}$.

We note that one can also obtain an understanding of how the spike correlations change when $N$ approaches the limit $1+\frac{1}{\hat{w}}$ by directly considering the non-expanded interdependence term of the correlation function, $\tilde{C}^1_{i\neq j}(\omega)$, cf. Eqs. 4,S.1,S.10. For this, we first set $(N-1)\hat{w} = 1-\epsilon$ with $\epsilon > 0$. Eq. S.10 becomes

$$\begin{aligned}\tilde{C}^1_{i\neq j}(\omega) = r\Bigg(&\frac{\hat{w}(1+i\omega\tau_\mathrm{s})(1-i\omega\tau_\mathrm{s})}{(1+i\omega\tau_\mathrm{s}+\hat{w})(1-i\omega\tau_\mathrm{s}+\hat{w})(\epsilon-i\omega\tau_\mathrm{s})}+\\ &+\frac{\hat{w}(1-i\omega\tau_\mathrm{s})(1+i\omega\tau_\mathrm{s})}{(1-i\omega\tau_\mathrm{s}+\hat{w})(1+i\omega\tau_\mathrm{s}+\hat{w})(\epsilon+i\omega\tau_\mathrm{s})}+\\ &+\frac{N\hat{w}^2(1+i\omega\tau_\mathrm{s})(1-i\omega\tau_\mathrm{s})}{(1+i\omega\tau_\mathrm{s}+\hat{w})(\epsilon+i\omega\tau_\mathrm{s})(1-i\omega\tau_\mathrm{s}+\hat{w})(\epsilon-i\omega\tau_\mathrm{s})}\Bigg).\end{aligned} \tag{S.34}$$

In our models we have $\hat{w} \ll 1$ (cf. Table 1). In particular, taking the limit $N \to 1+\frac{1}{\hat{w}}$ implies $N \gg 1$. We can thus additionally use $\hat{w} \approx (1-\epsilon)/N$. Eq. S.34 then simplifies to

$$\tilde{C}^1_{i\neq j}(\omega) \approx r\frac{(1-\epsilon)(1+\epsilon)}{N(\epsilon^2+\omega^2\tau_\mathrm{s}^2)}. \tag{S.35}$$

The limit $N \to 1+\frac{1}{\hat{w}}$ further implies that $\epsilon \ll 1$, which leads to

$$\tilde{C}^1_{i\neq j}(\omega) \approx \frac{r}{N}\frac{1}{\epsilon^2+\omega^2\tau_\mathrm{s}^2}. \tag{S.36}$$

Taking the inverse Fourier transform, we obtain in the time domain

$$C^1_{i\neq j}(t) \approx \frac{r}{2\tau_\mathrm{s}\epsilon N}\exp\left(-\frac{\epsilon\,|t|}{\tau_\mathrm{s}}\right). \tag{S.37}$$

Thus, $C(t)$ and hence the distribution of time lags of spike pairs become flatter and flatter as $\epsilon \to 0$. Inserting Eq. S.37 into Eq. 7 yields the weight changes

$$\overline{\Delta W_{ij}}(N) = \int_{-\infty}^{\infty} \mathrm{d}t \; F(t)(C_{ij}^{0}(t) + C_{ij}^{1}(t)) \tag{S.38}$$

$$\approx f_0 r^2 + \frac{r}{2\tau_\mathrm{s}\epsilon N} \int_{-\infty}^{\infty} \mathrm{d}t \; F(t) \exp\left(-\frac{\epsilon\,|t|}{\tau_\mathrm{s}}\right) \tag{S.39}$$

$$\approx f_0 \left(1 + \frac{1}{2\lambda_0 \tau_\mathrm{s} N}\right) \frac{\lambda_0^2}{\epsilon^2}. \tag{S.40}$$

In the last line we used Eq. 14 and that $\exp\left(-\frac{\epsilon|t|}{\tau_\mathrm{s}}\right) \approx 1$ in the relevant range of $t$, where $F(t)$ is noticeably different from zero. Eq. S.40 is consistent with main text Eq. 20, which yields the same result after setting $1 + \hat{w} \approx 1$, substituting $(N-1)\hat{w} = 1 - \epsilon$ and using $\epsilon \ll 1$.

### S.3 Lower bound of the input strength for the storage of a new assembly

To analytically estimate the strength of feedforward input that is required to embed an assembly, we consider a network where the recurrent weight strength is initially zero and focus on a pair of neurons of the future assembly. The common external input evokes correlated spiking in both neurons, which leads to a change of the recurrent weights between them. The common external input thereby acts like a common presynaptic network neuron with rate $r_{\text{in}}$ and projection strength $w_{\text{in}}$ to both neurons. We can thus use Eq. 11 to compute the weight change $\overline{\Delta W_{ij}}$. The external input increases the spike rate of the considered neurons from the spontaneous rate $\lambda_0$ to the rate $\lambda_0 + r_{\text{in}}w_{\text{in}}$, which causes a rate-based weight change of $f_0 r_i r_j$. Further, we have spike correlations in the two neurons, which are due to a chain of $\alpha = 1$ and $\beta = 1$ connections from a spiking source neuron (the external input) with presynaptic rate $r_{\text{in}}$ (the external input projects directly on both considered neurons). Since we assume that there are no recurrent weights in the network, the spikes cannot propagate and there are no further motifs that evoke correlated spiking. Eq. 11 thus yields

$$\overline{\Delta W_{ij}} = f_0(\lambda_0 + r_{\text{in}}w_{\text{in}})^2 + r_{\text{in}}f_{11}w_{\text{in}}^2. \tag{S.41}$$

Assembly creation requires that there is net potentiation $\overline{\Delta W_{ij}} > 0$ of the weights between the considered neurons. For given $r_{\text{in}}$ Eq. S.41 provides a condition on $w_{\text{in}}$,

$$w_{\text{in}} > \frac{-2f_0\lambda_0 r_{\text{in}} + \sqrt{(2f_0^2\lambda_0^2 r_{\text{in}}^2 - 4f_0\lambda_0^2(r_{\text{in}}^2 f_0 + r_{\text{in}}f_{11})}}{2r_{\text{in}}^2 f_0 + r_{\text{in}}f_{11}} =: w_\theta. \tag{S.42}$$

To numerically check and further investigate this result, we plot the summed interconnecting weights of a set of targeted neurons after stimulation in Fig. S4. We find that assembly formation, reflected by an increase of the connections between the stimulated neurons, indeed starts for $w_{\text{in}} \approx w_\theta$. Due to the inherent stochasticity of the networks, there is already an increase in the connection weights for $w_{\text{in}}$ below $w_\theta$. The weight plasticity noise becomes stronger with larger learning rate. Indeed we observe a less pronounced kink in the relationship between the summed weights and $w_{\text{in}}$ at $w_{\text{in}} \approx w_\theta$ for larger learning rates. In Fig. S4 we keep the product of learning rate and input duration constant, such that the impact of constant drifting proportional to $\mu$ stays the same. This is, however, not necessary for the observation.

### S.4 Parameters

| | $N$ | $\hat{w}$ | $\tau_{\mathrm{s}}$ | $A_{\mathrm{p}}$ | $A_{\mathrm{d}}$ | $\tau_{\mathrm{p}}$ | $\tau_{\mathrm{d}}$ | $\mu$ | $\lambda_0$ |
|---|---|---|---|---|---|---|---|---|---|
| Fig. S1 | | | 0.01 s | 0.08 | -0.053 | 0.025 s | 0.05 s | | |
| Fig. S2 | | | 0.01 s | 0.08 | -0.053 | 0.025 s | 0.05 s | | 0.15 Hz |
| Fig. S3 | | | 0.01 s | 0.08 | -0.053 | 0.025 s | 0.05 s | | 0.15 Hz |
| Fig. S4 | 50 | 0.026 | 0.01 s | 0.08 | -0.066 | 0.035 s | 0.05 s | 0.07 | 0.15 Hz |
| Fig. S5 | 60 | 0.024 | 0.01 s | 0.08 | -0.053 | 0.025 s | 0.05 s | 0.045 | 0.15 Hz |
| Fig. S6 | 80 | 0.024 | 0.01 s | 0.08 | -0.053 | 0.025 s | 0.05 s | 0.055 | 0.15 Hz |
| Fig. S7 | 72 | 0.056 | 0.01 s | 0.08 | -0.053 | 0.025 s | 0.05 s | 0.148 | 0.2 Hz |

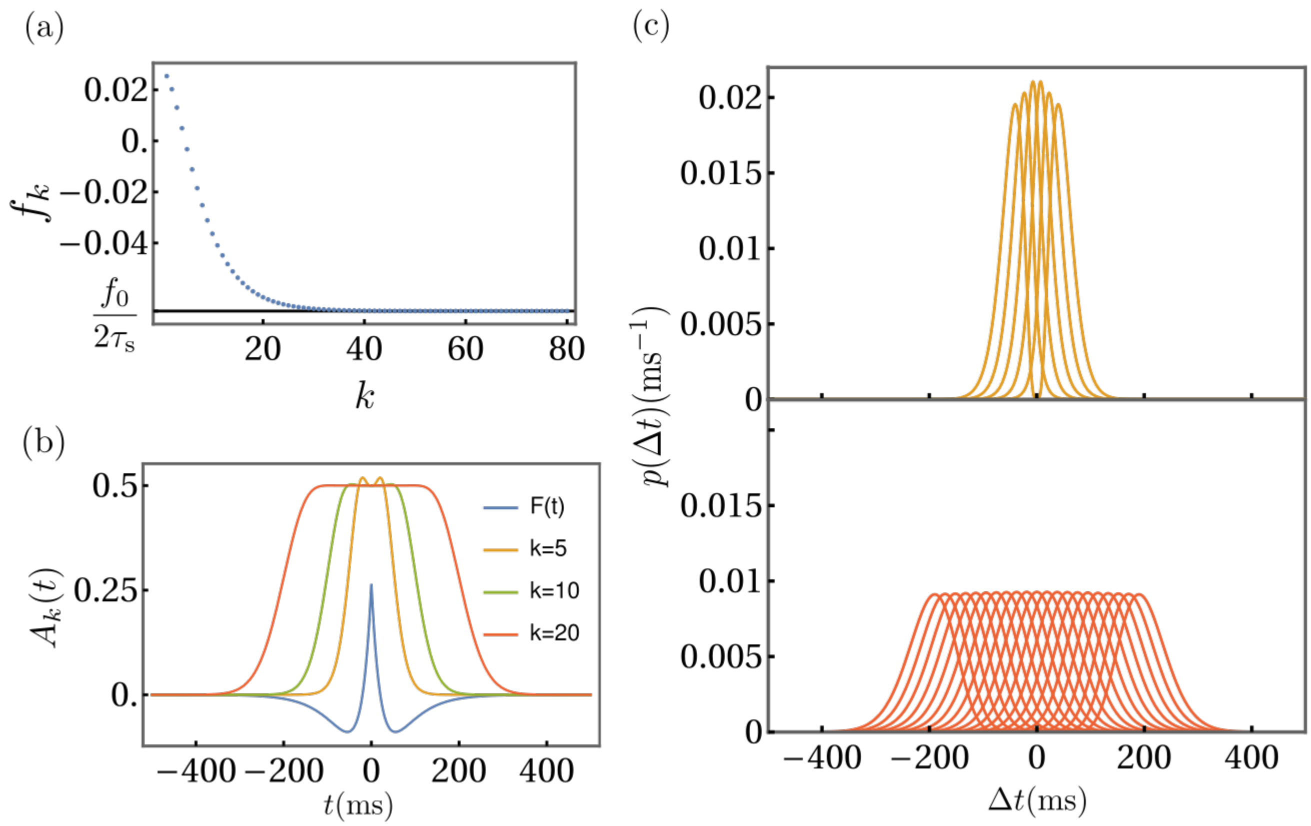


Figure S1: Asymptotic behavior of sums $f_k$ of motif contributions. (a) Convergence of $f_k$ to $\frac{f_0}{2\tau_s}$ for high orders of $k$. (b) Sums of convolutions of the synaptic current functions $A_k(t)$ (see Eq. S.31), for different values of $k$. As discussed in section S.2, these curves can be interpreted as resulting from the superposition of distributions of time-lags for spike cascades of total length $k$ affecting pre- and postsynaptic neurons (see (c)). For high orders, $A_k(t)$ becomes a plateau of height $\frac{1}{2}$. The learning window (blue, a.u.), is shown to illustrate the time-scales. (c) Probability distributions of the time lags $\Delta t$ between the last spikes of two spike-cascades of different lengths $\beta$ and $\alpha$. The total cascade length $k = \alpha + \beta$ equals 5 spikes (upper subpanel) or 20 spikes (lower subpanel). For larger $k$ the distributions have larger variance and are more spread out. Their superpositions therefore assume the widening plateau shapes displayed in (b).

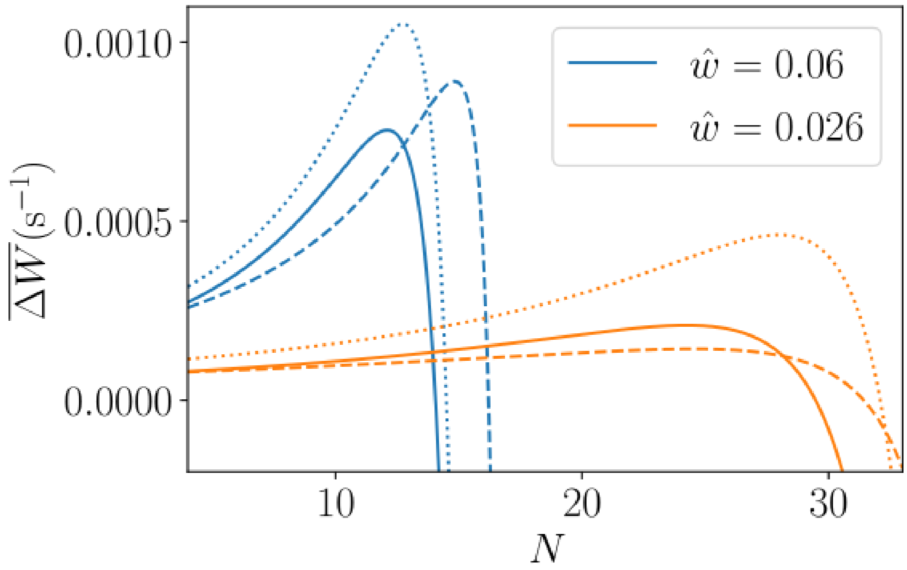


Figure S2: Comparison of the time-averaged weight change within homogeneous assemblies as in Fig. 3a, for full network plasticity (Eq. 15, solid lines), for truncated plasticity without the impact of higher order network motifs (we keep only the terms proportional to $f_0$, $f_{10}$, $f_{01}$ and $f_{11}$ in Eq. 11, dashed lines) and for full plasticity without the rate-rate interaction term (we omit the term proportional to $f_0$ in Eq. 11, dotted lines). The figure shows results for two values of $\hat{w}$, which were also used in Fig. 3a.

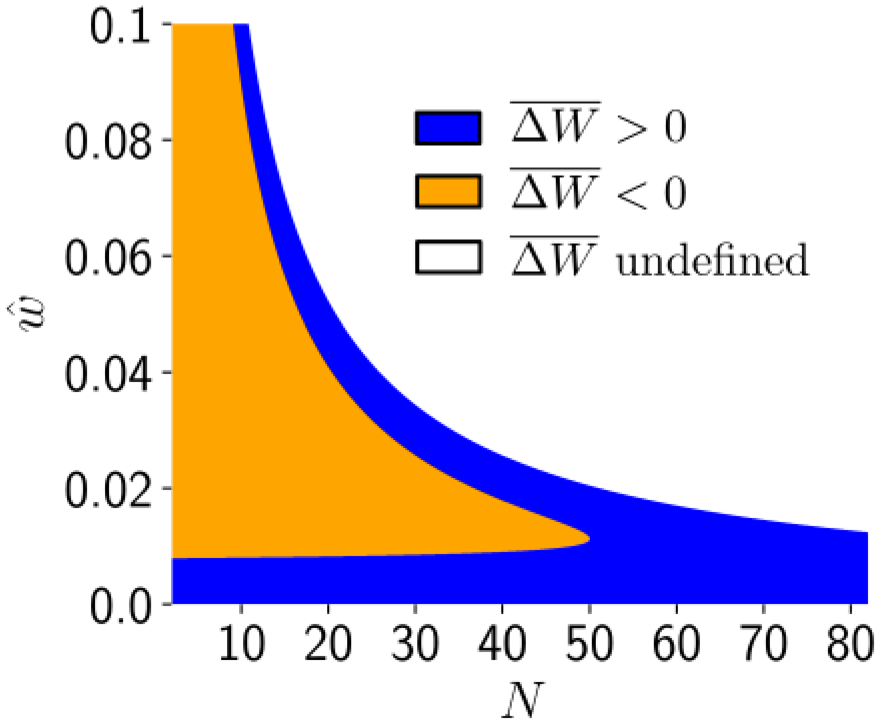


Figure S3: Phase diagram of the time-averaged weight change in fully connected assemblies, with respect to assembly size $N$ and assembly weights $\hat{w}$. The changes are obtained from main text Eq. 15. In the orange area the weights increase and in the blue area the weights decrease. In the white area the activity is divergent.

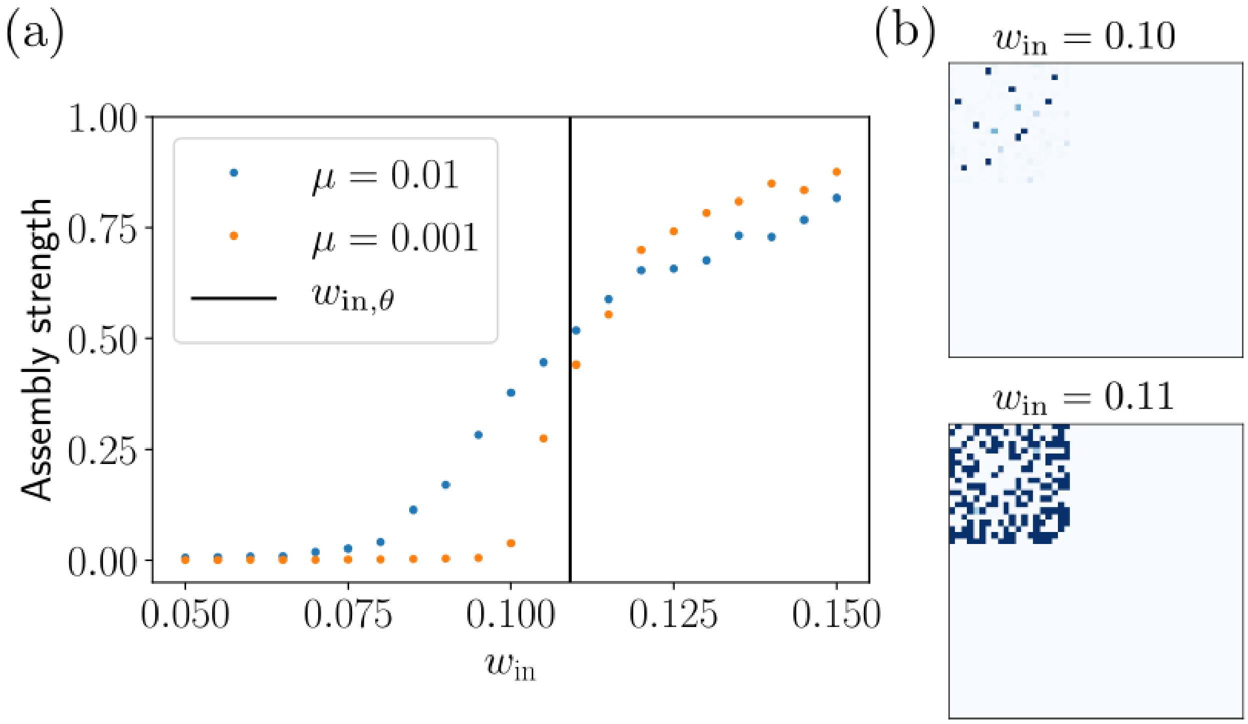


Figure S4: Lower bound of the input strength for the storage of a new assembly. (a): Summed weights between the target neurons relative to the maximum summed strength $N(N-1)\hat{w}$, after correlated input with different input strengths $w_{\text{in}}$, for two different values of $\mu$ and durations $T$ such that $\mu T = 1 \times 10^3\,\text{s}$. The theoretical threshold for assembly formation $w_{\text{in},\theta}$ is highlighted by a black line. (b): Final weight matrices from simulations in (a) for $\mu = 0.01$ and two values of $w_{\text{in}}$.

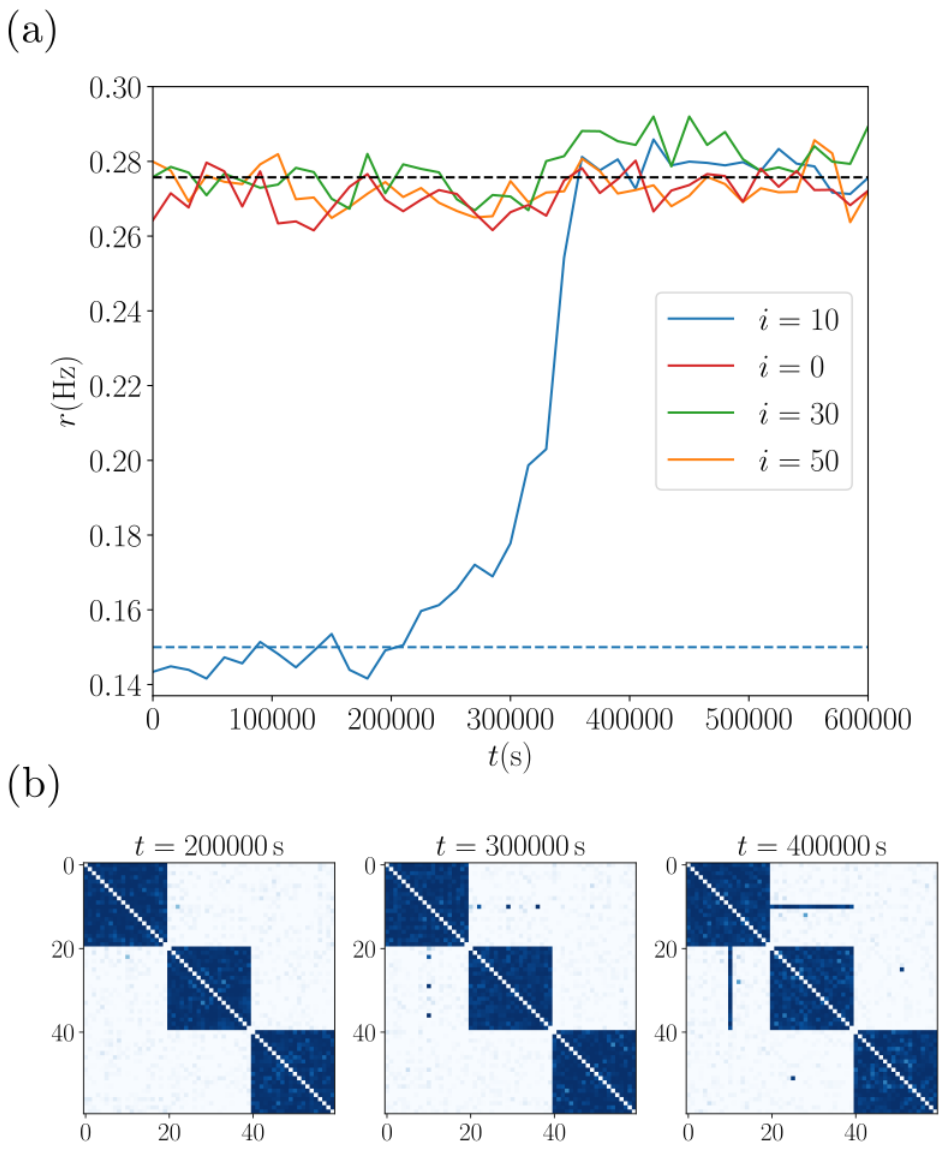


Figure S5: Firing rate evolution of a neuron that spontaneously joins an additional assembly. (a) The neuron with index $i = 10$ has a spontaneous rate $\lambda_{x,0}$ ($x = i = 10$) given by Eq. 28. Its time-averaged firing rate while connected to only one assembly is thus approximately given by $r \approx \lambda_{x,0} + \lambda_0/(1-(N-1)\hat{w}) - \lambda_0$ (blue dashed line). After it has joined another assembly (at around 350000s, see (b)), its rate is approximately given by Eq. 14 (black dashed line), i.e. it is approximately equal to that of other assembly neurons. For comparison, sample rates of three neurons are shown: of neuron 0 from assembly 1 (the assembly that the overlap neuron is originally in), of neuron 30 from assembly 2 (the assembly that the overlap neuron joins) and of neuron 50 from assembly 3 (the non-overlapping assembly). The rate of neuron 0 is initially slightly lower due to the smaller rate of neuron 10. The rate of neuron 30 increases after overlap formation since the second assembly then has an additional neuron. (b) Weight matrices of the network at different times.

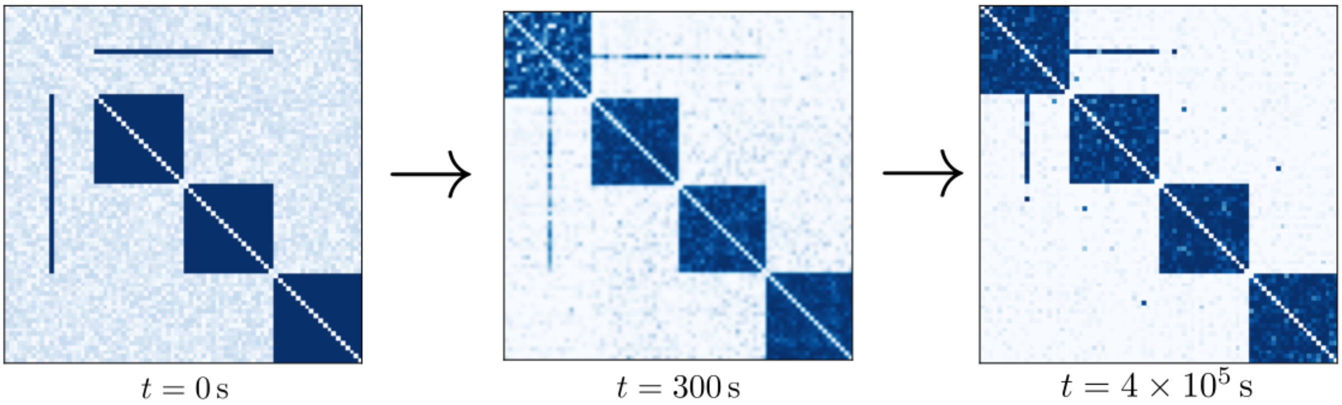


Figure S6: A network with preexisting assemblies, background neurons and a neuron that is part of two assemblies, learns a new assembly. The figure displays the weight matrices of the initial network configuration (left hand side), after stimulation (middle) and after a longer time (right hand side). The stimulation of neurons 1-20 lasts for 300 s with the same stimulation protocol as in Fig. 4. The initially strong connections of the "overlap neuron" (neuron 10) with two other assemblies experience depression after the neuron is recruited to the new assembly (middle). Subsequently the neuron loses its connections to one of the assemblies it previously belonged to (right hand side).

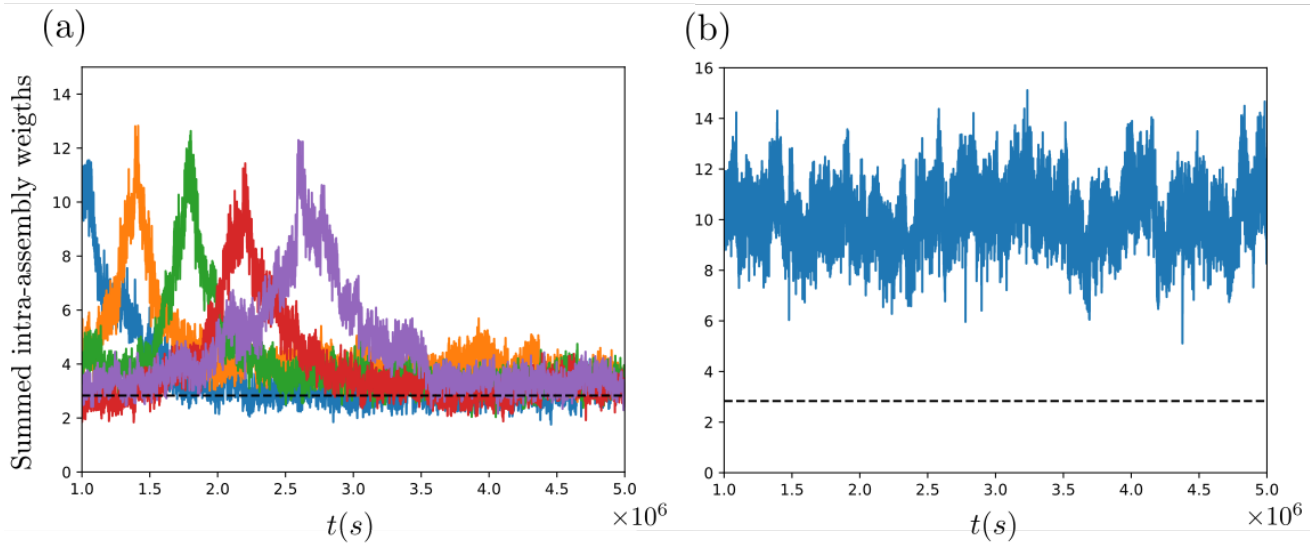


Figure S7: Drifting assembly dynamics. (a) The network with drifting assemblies of Fig. 8a is simulated over a longer time. We choose at five time points ($t = 1.0 \times 10^6$ s, $1.4 \times 10^6$ s, $1.8 \times 10^6$ s, $2.2 \times 10^6$ s, $2.6 \times 10^6$ s) an assembly as a reference and note the neurons forming it. Thereafter, we track the sum of weights between these neurons in the present, future and past weight matrices. For sufficiently long temporal distances this sum approximately reaches chance level (dashed black line, for an average size assembly), which we define as the average of the sum of interconnections in groups of randomly chosen neurons. We can therefore conclude that assemblies indeed completely drift, that is, there is no constant "stable core" set of neurons in an assembly. (b) Tracking of a single drifting assembly over time. At $t_0 = 1 \times 10^6$ s we choose assembly 1 (of the four assemblies) and set $t = t_0$. We then compute the sum of the weights at time $t + \Delta t$ ($\Delta t = 2$ s) between the neurons that formed assembly 1 at time $t$, to see if they are still strongly connected. We repeat the procedure using the actual assembly 1 at the new time $t = t_0 + \Delta t$ (which may have exchanged individual neurons compared to $t_0$). In particular, we again compute the weights between its neurons at $t + \Delta t$ and so on. We find that the sum of weights stays at a consistently high level. In other words, neurons that form assembly 1 at a time $t$ are still strongly connected at $t + \Delta t$. The change of the assembly is therefore gradual; most neurons that are part of the assembly at $t$ are still part of it at $t + \Delta t$. This implies that the assembly can be tracked over time, despite the complete drift.
